# NeMu: A Comprehensive Pipeline for Accurate Reconstruction of Neutral Mutation Spectra from Evolutionary Data

**DOI:** 10.1101/2023.12.13.571433

**Authors:** Bogdan Efimenko, Konstantin Popadin, Konstantin Gunbin

## Abstract

One of the most important characteristics of each contemporary model of molecular evolution is the assumption that mutations occur in a constant manner; however, in the real world, the mutations are determined by the combination of the effects of DNA replication and repair. This affects the nucleotide composition of the genome and guides not just neutral but adaptive evolution^1^. Mutation accumulation experiments are the de facto standard for the neutral mutation spectra estimation. However, recent studies have demonstrated that the mutation fraction under selection pressure is significantly underestimated in mutation accumulation experiments, and, therefore the precise extraction of neutral mutation spectra from mutation accumulation experiments is not trivial^2^. To unravel the neutral mutation spectra, it is very important to analyze all the mutations available in depth, based on the evolutionary timescale, taking into consideration all the existing knowledge. In order to facilitate this analysis, we have created a novel pipeline, called NeMu (https://biopipelines.kantiana.ru/nemu/).

## INTRODUCTION

To rephrase the Nobel Lecture of Sydney Brenner, many important things in science are drowning in an ocean of data and unfortunately starving for discovery (Brenner, S. (2003). NOBEL LECTURE: Nature’s Gift to Science. Biosci. Rep. 23:225–237^3^). For example, more and more data about human evolution reveal that evolutionary models with constant mutation rate inadequately describe how variation accumulates both for inter-^4,5^ and intra-specific^6–8^ time scales. On the other hand, It is generally accepted that because of the different agents of DNA damage and failures in the DNA repair process, many cancers exhibit somatic hypermutability in the certain triplet patterns^9,10^. Thus, in the real world, the mutations are not random and constant during time, but are determined by the combination of DNA replication and repair that control the species’ and populations’ overall mutation spectra.

The specificity of the mutation process affects the nucleotide composition of the genome and guides not only neutral but adaptive evolution^1^. For example, systematic studies of the mutational spectra in various species such as *Escherichia coli*^1,11,12^; yeast^13^; *Mycobacterium smegmatis*^14^; *Drosophila melanogaster*^15^; *Arabidopsis thaliana*^16^; human^17^; influenza A viruses^18^; SARS-CoV-2 viruses^19^) have clearly demonstrated the presence of nucleotide substitution biases, including the well-known transition bias, the CpG bias and other nucleotide context-dependent biases. The observed fixations can be predicted as a function of the mutation spectra, e.g., the amino acid fixations in the proteomes^12,20^ between two closely related species or the dissimilarity of regulatory genetic elements in closely related genomes^21^. It is of importance, that a huge over-representation of the transitions (for example changes from G to A) over the transversions (for example changes from G to C) is not compatible with the fact that the transversions are not clearly more conserved than the transitions in the context of the interchangeability of amino acids^22^. However, the shift to transitions usually shapes parallelism in adaptations between species^23,24^. Ultimately speaking, biased mutagenesis (or biased DNA repair) shapes genomic characteristics, drastically orienting evolutionary pathways for both neutral substitutions and even adaptive fixations^12^.

Mutation accumulation experiments^25^ are the de facto standard for the neutral mutation spectra estimation, yet. However, this type of experiment is time-consuming, expensive and usually requires analysis of only well-studied organisms^11,13^. It is generally accepted that natural selection plays a minimal role in the mutation accumulation experiments. However, recent studies have shown that the beneficial mutation fraction is greatly underestimated in mutation accumulation experiments^2^, and, for this reason, in general, the precise extraction of neutral mutation spectra from mutation accumulation experiments is not a trivial procedure^2,26^.

An alternative and more straightforward way to estimate neutral mutation spectra is to analyze natural intraspecific polymorphism data^14,17^ within the context of intraspecific evolution and natural selection. This approach to estimating neutral (or nearly neutral) mutation spectra is less confused by uncontrollable experimental conditions. Thus, to disentangle neutral (or near neutral) mutation spectra from selected spectra, it is very important to thoroughly analyze all available mutations, based not only on a single time point (e.g. in the current evolutionary state), but also on the evolutionary time scale, taking into consideration all the available knowledge such as: 1) the theoretical level of various forms of mutation affecting fitness (e.g., synonymous, non-synonymous, four-fold degenerative synonymous, etc.), 2) the potential selection constraints and/or signatures of natural selection, 3) mutability biases, the specificity of their species/population characteristics or, on the other hand, their reproducibility, 4) the variability and conservation of single nucleotide compositions and DNA motif compositions, 5) the rigor of reversibilities (or non-reversibilities) of nucleotide fixation rate models on different phylogenetic lineages, etc. To do so, we first have to perform numerous data quality tests to ensure that our inferences are not based on errors or confounding factors. To make it easier to study molecular mutation spectra and to accurately estimate the neutral mutation spectrum component, we created an automated and user-friendly pipeline called NeMu. This is, to the best of our knowledge, the first software in this field so far.

## RESULTS & DISCUSSION

### (1) Intuitive and Advanced interface for rapid reconstruction of mutational spectra for any gene of any species

We have developed a user-friendly interface for our NeMu pipeline, we created simple and handy (interactive) web-pipeline, which enables non-bioinformaticians to obtain mutational spectrum and to disclose the neutral component of this mutation spectrum for any gene of any species in just minutes. The pipeline offers various input options, with the two basic ones: a species name and a protein sequence. The pipeline also provides several types of outputs, with the basic output being the 12 substitution rates (referred to as the 12-component mutational spectra). This pipeline features high flexibility, from basic to highly advanced, with the ability to tune dozens of additional parameters to meet the specific needs. On Fig. 1 we show a schematic diagram of the NeMu pipeline being implemented.

**Fig. 1.**
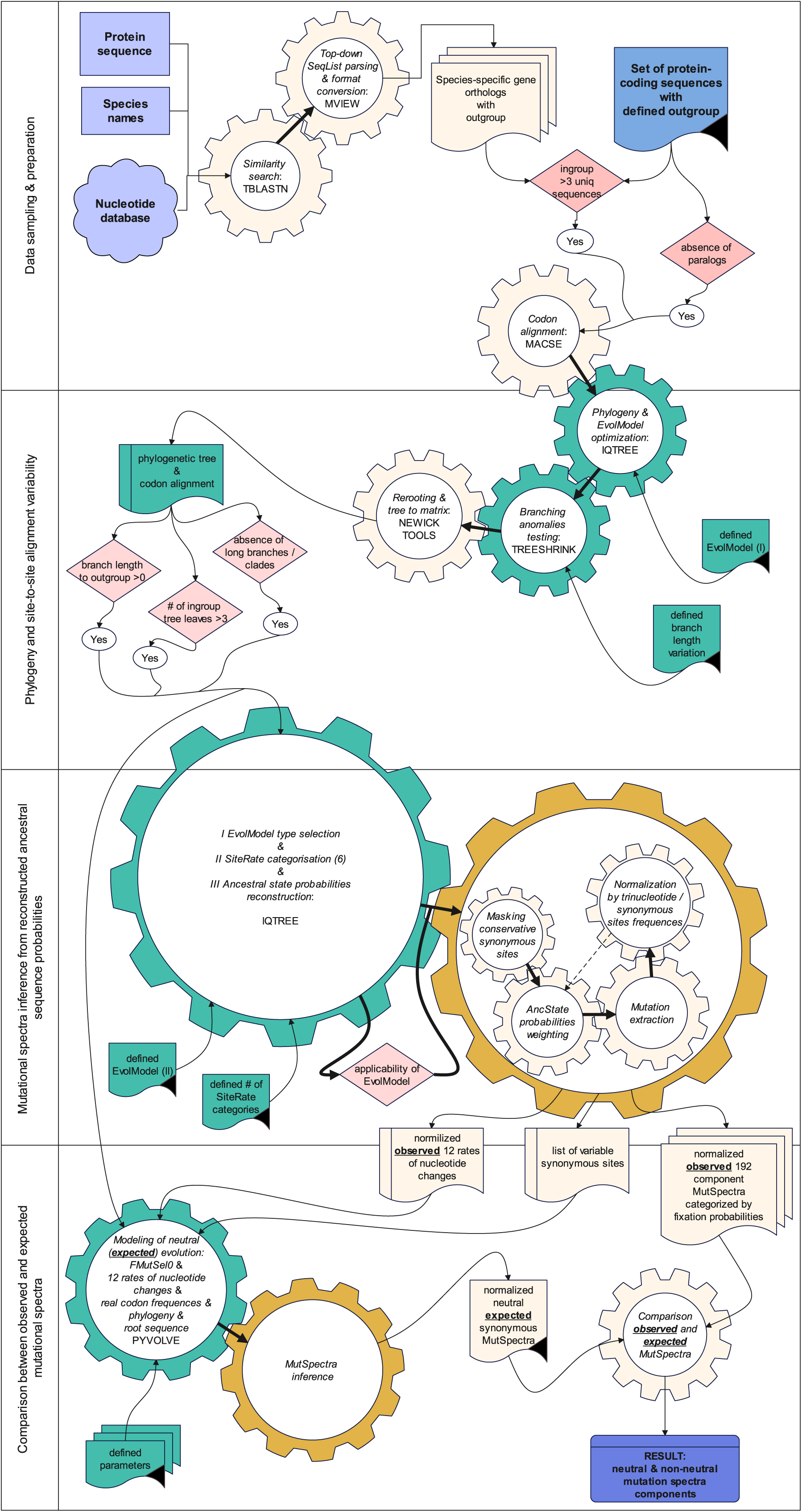
General scheme of the NeMu pipeline. The ivory color in the diagram indicates the automatic stages and substages, gold color indicates the automatic unified mutation spectra inference stage. The blue and violet colors in the diagram indicate start and end stages. The green color in the diagram indicates the stages that may require the researcher’s control, for example: 1) the choice of the nearest outgroup sometimes cannot be made automatically due to the possibility of convergent evolution; 2) the reconstruction of the phylogenetic tree using the maximum likelihood method requires the selection of the optimal Markov model for the evolution of protein-coding genes, often the automatic choice of single Markov models is difficult^59,81–83^; 3) the selecting out individual branches, subtrees, invariant or conservative sites of an ingroup (analyzed data set) is often associated with a dilemma in identifying the nature of the accelerated evolution of sequences (sequencing errors or real data^52,84^); 4) the reconstruction of the probability vectors of ancestral states (at each position of the multiple alignment and on each internal branch of the phylogenetic tree) can be complicated by non-optimal selection of Markov models of evolution for the ancestor reconstruction^47,48,85,86^. Internal checkpoints of NeMu pipeline, which stop pipeline execution in case of failure, are highlighted in red in the diagram: 1) a minimum number of sequences for the reconstruction of a stable phylogeny; 2) minimum number of terminal branches in an ingroup; 3) statistical difference between the lengths of ingroup branches and the length of outgroup branches (no errors in automatic tree re-rooting); 4) absence of natural selection signatures (e.g., sequence evolutionary conservation); 5) applicability of the time-reversible Markov model of sequence evolution for constructing ancestral sequences^47,48,50^.

The general aim of our pipeline is to disclose the neutral mutation spectra. To do so, we based our estimations on synonymous nucleotide substitutions of protein coding genes. The majority of evolutionary studies indicate that this type of nucleotide substitution may be used as a surrogate for neutral (or nearly neutral) mutation spectra. However, there is considerable evidence that specific classes of synonymous nucleotide substitutions may be beneficial or harmful^27–29^. Thus, in order to be as accurate as possible in the transfer of synonymous substitutions to neutral estimates, to manage observations, we have implemented simulations along with numerous data filters: we have based simulations on mutation-selection model taking into consideration the codon fitness and the codon usage. Additionally, we carefully exclude any invariable sites from the estimates.

Despite the great flexibility of the pipeline and the many options available, we construct an intuitive graphical user interface https://biopipelines.kantiana.ru/nemu/. Genes encoded in the mitochondrial genome can be processed automatically, and manually prepared samples of genes can be processed if they are encoded in nuclear DNA. Fig. 2 demonstrates the implemented interface of the NeMu pipeline. Execution of the NeMu pipeline requires several main and optional input arguments (see NeMu Manual; https://github.com/mitoclub/nemu-pipeline/wiki). To ensure user simplicity and pipeline flexibility, we have implemented two main input methods: automatic (we use MIDORI2^30^ mtDNA database for sequence retrieval) and manual sampling. To run the pipeline through the automatic data retrieval method, users are required to select mode “Auto”, upload a sequence fasta file with single amino acid sequence, paste species name, select genetic code and gene name for specific tblastn search. To run the pipeline through the manual method of sample construction, the user is required to select mode “Manual”, select level of analysis (species-specific or comparative-species) and paste the name of the outgroup sequence that must be present in the aligned (preferred but not required) fasta file with multiple nucleotide sequences. In addition, there are also optional parameters of several processes. For instance, users can use a preferred substitution model used by IQTREE in the tree building and ancestral states reconstruction steps, turn off “Use probabilities” mode to consider the most probable nucleotide in each site instead of nucleotide probability distribution, control mutational spectra inference, for example, derive spectrum for fourfold synonymous mutations, for internal or terminal tree branches and so on. Advanced users can modify these and other parameters depending on their own needs. After the pipeline execution is completed, the user is able to access the output page, which provides the option to download the results within a time period of 7 days. Additionally, users who have specified an email address will receive a notification with the output archive file attached. There are multiple directories with multiple output files stored inside the output archive: tables with mutations and mutation spectra in the tables directory, spectra barplots in the images directory etc. Additionally, there are plenty of intermediate outputs available to control the execution of each process. These include log files for all crucial pipeline steps, a phylogenetic tree file and a site rates table. As a result, users are able to check and rerun different steps manually if they want to use their own custom settings.

**Fig 2.**
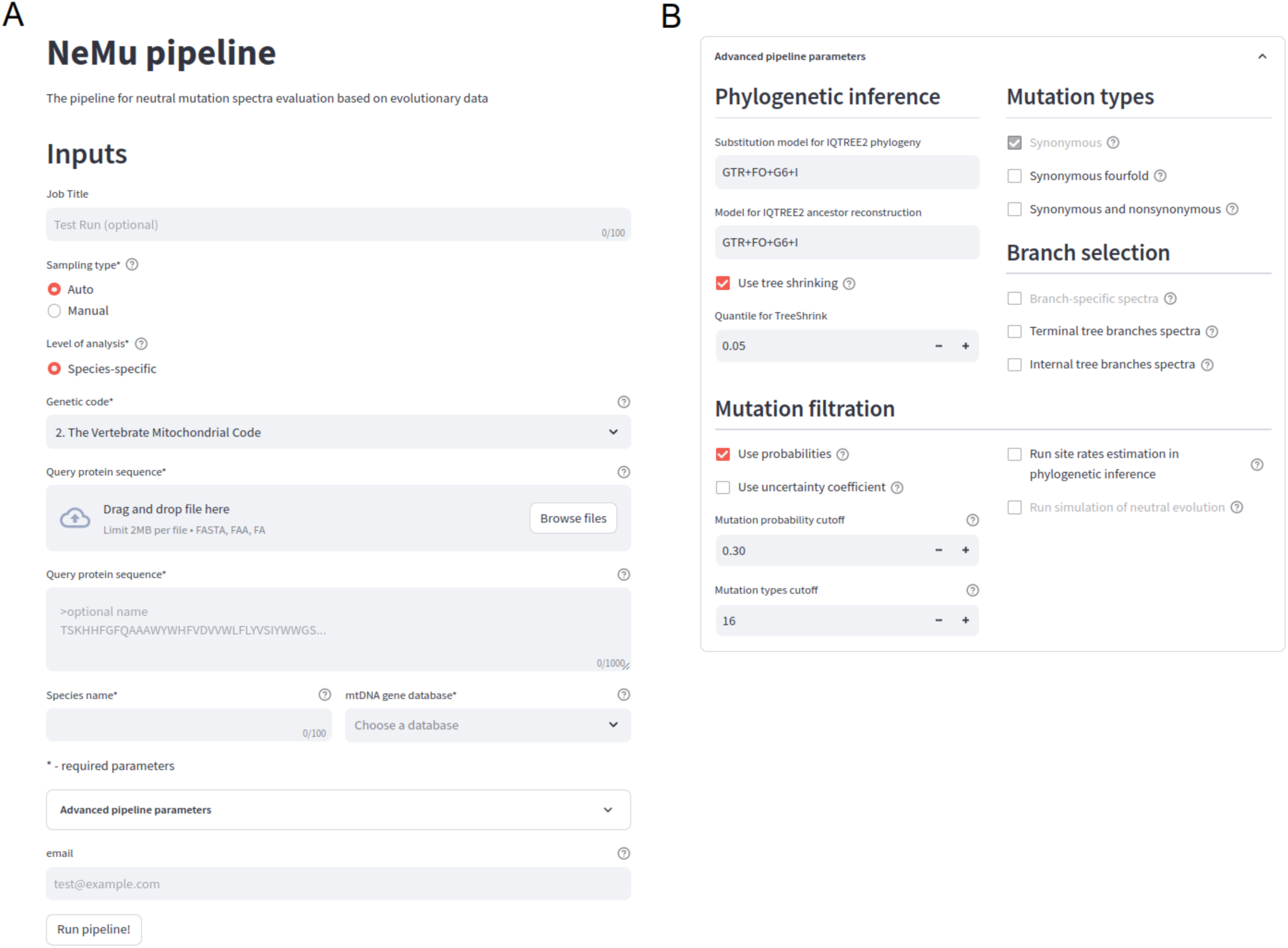
The interactive intuitive graphical user interface of the NeMu pipeline. (A) The main page of the NeMu pipeline web-interface. (B) Advanced pipeline parameters interface.

The advanced users are welcome to run the NeMu pipeline in an isolated stationary environment on different operating systems through the Singularity^31^ container. Definition file of the NeMu pipeline container can be found in the Github repository (https://github.com/mitoclub/nemu-pipeline/tree/master/singularity). It contains all programs and scripts for running a pipeline on all operating systems with Singularity installed. For instance, the NeMu pipeline in the interface utilizes this container by default during execution. A convenient way of using the NeMu pipeline on high performance clusters is headless usage mode, e.g. by running separate steps of the NeMu Singularity container. Thanks to the Nextflow scripting language^32^ that represents pipelines as collections of independent processes, each process in our NeMu pipeline can perform one or more commands or scripts on various programming languages (Bash, Perl, Python, Java, various binaries compiled from C and CPP sources). Therefore, for the advanced users, we have provided a convenient way to modify the pipeline backbone (Nextflow scripts) to any purpose.

### (2) Data sampling and data preparation

The primary objective of our NeMu pipeline is to extract the neutral (or near neutral) component of the species-specific mutational spectrum. In the pipeline, it may be reached by two alternative methods of sample construction: 1) by sequential automatic extraction of orthologous mitochondrial DNA sequences encoding proteins for each species from a user-defined sets of gene names and species names using MIDORI2^30^ mitochondrial database (sequential automatic data retrieval method) or 2) giving the predefined set of protein-coding orthologous DNA sequences (it is preferred to align sequences in multi-fasta file, but it is not mandatory) into the pipeline (the manual method). Both sample construction methods require protein coding sequences with clear orthology. We require that because the alignment consisting of a mix of orthologs and paralogs has a significant probability of fixations (even synonymous) under positive selection due to the functional divergence of paralogs^33–38^.

Sequential automatic data retrieval method of sample construction enables the user to automatically construct a set of mitochondrial neutral mutational species-specific spectrum as a result, however, the completeness of each of these spectra depends on the number of substitutions observed in the species-specific dataset. Based on a simple requirement of any phylogenetic tree resolution, we set the minimum number of unique (nonredundant) sequences in the species-specific ingroup to 4. Nevertheless it is not sufficient to construct complete mutational spectra with 192 components (12 possible directions of simple nucleotide substitutions multiplied by 4^2^ combinations of two neighboring nucleotides), especially if these four sequences have a low number of synonymous substitutions. Thus, for this method of sample construction, it is very important to check the compromise between the number of included species and the exhaustiveness of species-specific spectra. It is important to note that this sample construction method have a simple technical insight enabled us to be sure that in the entry we still have one-to-one strict orthologs: the list of retrieved mitochondrial sequences (which are unique/non-duplicated in mtDNA) is automatically analyzed from top, best homologous to query, to bottom, last homologous to query, and analysis stops when the sequence belongs to other species found (this sequence used as an outgroup).

The manual method of sample construction (e.g., set of aligned protein coding sequences as input) allows the user to build a single neutral (or nearly neutral) mutational spectrum specific to non-species taxon (e.g., the taxon rank of family or order); in addition, both mtDNA genes and nuclear DNA genes can be utilized to create the sequence sample in this manner. If possible, it is recommended that the sequence set be filled in by strict orthologs in this sample construction method. We cannot recommend this method of sample construction to a higher taxonomic rank (e.g., the rank of taxonomic class or type) because of the much greater ambiguity in the classification of neutral (or nearly neutral) alignment positions with synonymous fixations and because of the synonymous substitution saturation^39,40^. To construct a mutational spectrum for higher taxonomic rank we recommend procedures described in our analysis of mtDNA mutation spectra of *chordata* species^41^. The manual method of sample construction is often exacerbated by unnoticed and not trivial difficulties: it is important for the user to make sure that the pairwise distance distribution between the aligned sequences is uniform (this means that it should not be multimodal), otherwise, the mutations on the longest tree branches, which correspond to outlayer branch distances, could cause a significant bias in the mutational spectrum^42^. The general solution of this problem we provided in the next, phylogenetic, stage of data processing. However, we recommend the users to pre-check the dataset they upload to avoid automatic removal of sequences from the sample.

Following the construction of the sample, in the pipeline we align the protein coding sequences of the sample sequence set by MAFFT v7.505^43,44^, which is utilized to quickly align translated protein sequences and then translate this alignment to codons using reportGapsAA2NT MACSE module. Also, we allow advanced users to enable MACSE v2.06 aligner^45^ instead of MAFFT to run codon alignments directly.

### (3) Phylogeny and site-to-site alignment variability

Next steps for quality control prior to mutational spectra reconstruction are 1) the reconstruction of the phylogeny that describes the relations between all the sequences of a sample, 2) re-rooting of phylogeny, which is necessary for normal linear orientation of phylogenetic relationships over time (especially important if non-reversible models of evolution are preferred), 3) detecting branching anomalies, e.g. long branches in a phylogenetic tree and/or close to zero outgroup branch length, and 4) masking of conservative alignment sites for subsequent reconstruction of mutational spectra.

For phylogenetic reconstruction we used IQTREE 2.2.0^46^. We chose this software because of the fast computation speed and flexibility in the evolutionary model setting. This software has made it possible to simply use and test the applicability of time reversible and also non-reversible evolutionary Markov models for data analysis^47,48^. In the case of very time limited evolution (species-specific analysis for non-model organisms having very limited number of closely related sequences in the dataset), non-reversible Markov models are the only models that accurately reflect sequence evolution^49,50^.

The re-rooting of phylogeny and other classic manipulations of phylogeny topology is conducted by Newick tools 1.6.0 package^51^. We re-root the phylogenetic tree automatically when the user chooses the sequential automatic data retrieval method, while the option to manually re-root the tree by the user’s outgroup is available for the manual method of sample construction.

The pre-final step in quality control prior to mutational spectra reconstruction is to detect branching anomalies. If the sequential automatic data retrieval method has been chosen, we simply test the outgroup branch length for inequality to zero. If the length of the outgroup branch is zero, then outgroup is highly likely to belong to ingroup (due to, e.g., the error in sequence annotation in MIDORI2 database) or, otherwise, there’s a problem in the taxonomic definition of species, at least, based on the molecular data. If the user has selected the manual method of sample construction it is highly important to be sure that there are no obvious sub-trees or long branches in the reconstructed phylogeny (see above). To verify this, we use the TreeShrink software^52^ which implements an algorithm to find tree branches that could be removed to minimize the sum of branch lengths. TreeShrink is recommended by default for both sample construction methods because of the high frequency of various database errors (e.g., sequencing and/or annotation errors).

The final quality control step is masking of conservative sites for subsequent reconstruction of mutational spectra, including synonymous sites. To determine whether the alignment sites are possible subject to negative natural selection in the simplest case, it is enough to analyze the variation of the alignment sites. To do this, we use the variability class prediction of each alignment site based on the standard evaluation of rate heterogeneity (gamma distribution) across alignment sites, implemented in IQTREE 2.2.0^53,54^. For example, on Fig. 3A we show a rate heterogeneity characteristic for Hominida mtDNA genes. We obtained this estimate using both human data (GenBank database) and apes (Pan, Gorilla and Pongo, species listed in *Suppl. File 1*) data (GenBank database and ^55^) because of the likely ineffectiveness of substituting all possible neutral mutations over the course of human evolution (coalescence time of human mtDNAs including neanderthal and Denisovan mtDNAs is less than 1 MYA). Fig. 3B demonstrates a rate heterogeneity characteristic for Mus genus (*Mus musculus* data from GenBank database supplemented by other Mus species listed in *Suppl. 1*)

**Fig. 3.**
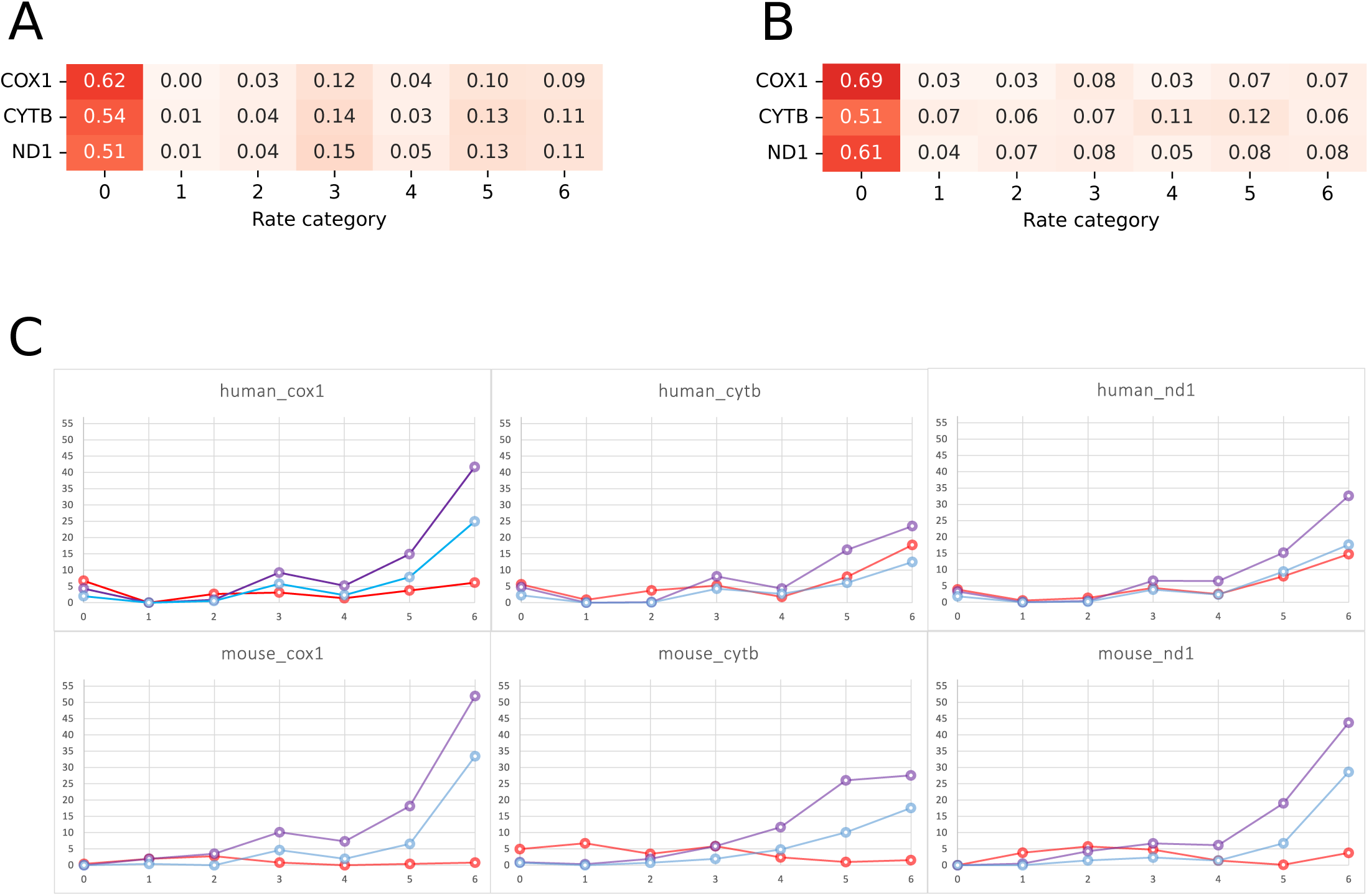
Variability of alignment sites in human and mouse mitochondrial genes. The fraction of alignment sites characterizing by 0 – 6 rate heterogeneity categories for 3 human (A) and 3 mouse (B) mitochondrial genes. Seven rate categories (from 0, totally conservative, to 6, totally variable) of gamma distribution were used in computations. (C) The percent (Y-axes) of substitution types and rate heterogeneity classes (X-axes) across alignment sites of human and mouse mitochondrial genes. Three substitution types were shown by different colors: total synonymous substitutions by purple, four-fold synonymous substitutions by blue, and non-synonymous substitutions by red.

### (4) Mutational spectra inference from reconstructed ancestral states probabilities: testing the validity of inferred mutational spectra

We have focused our main efforts on reconstructing the evolutionary history of DNA as accurately as possible. In doing so, first of all, we have a possibility for the user to exclude from consideration all mutations localized on the terminal branches of phylogenetic trees, because these mutations may originate from sequencing artifacts or some kind of harmful somatic events (the choice to exclude terminal branches from the analysis depends on the accuracy of the data is monitored by the user). Secondly, we have based our ancestral state reconstruction logic on the modified “uncertainty principle” proposed earlier to restore ancestral states^56^ on comparative species scale, which essence is the inability to evaluate simultaneously and precisely the rate of evolution of DNA sequences and ancestral states on a phylogenetic tree. In other words, the higher the rate of evolution on any branch of the phylogenetic tree is, the less likely it is to determine ancestry (and hence the types of nucleotide substitutions) at the extremities of that branch; and vice versa, the more the ancestral status is determined at the tree nodes, the less reliable the rate of evolution can be determined (i.e., the number of nucleotide substitutions per unit of time). It is very important, that the reconstruction procedures of ancestral states taking into account the “uncertainty principle” allows the selection of balanced evolutionary paths in terms of rates of evolution and definition of ancestral states (biologically meaningful or having sufficient information about the types and number of nucleotide substitutions; see illustrative Appendixes 1 and 2 to make it easier to understand). For higher (non-species taxon) taxonomic rank mutation spectra analysis we have suggested and implemented into our NeMu pipeline such an approach based on (see illustrative Appendixes 1 and 2 to make it easier to understand): 1) consideration of the probabilities of all alternative ancestral nucleotides at each multiple alignment position on each inner node of the phylogenetic tree; and 2) additionally (optional), consideration of the depth of the branches of a given phylogenetic tree by multiplying the probabilities of ancestral states of node *i* by the *y_i_ /d* ratio, where *y_i_*is the distance from node *i* to closest leaf (sum of branch lengths in the path from leaf to node *i*)*, d* is the geometric mean of distribution of distances from ingroup root to all leaves. Our algorithm is based on an exhaustive analysis of the probabilities of all ancestral states obtained at each node of a given phylogenetic tree using IQTREE 2.2.0^50,57–59^. These probabilities can be calculated from non-time-reversible Markov models that can significantly improve the final results^47,48^.

To test the effectiveness of our “uncertainty principle” normalization approach to filtering out errors/inaccuracies in ancestral reconstruction, on Fig. 4 and Fig. S2 we show pairwise comparisons (by cosine similarities) of mutation spectra obtained by various approaches on identical branches of mammalian ND1 and CYTB trees (the short data description of these datasets is given in Fig. S1). We compared our fast approach (named “uncertainty principle” normalization) with 1) the best but much slower (∼1000 times slower) analogous software available for general ancestral reconstruction, the PASTML software^60^, which allows, on the basis of combination of marginal posterior probabilities approximation (MPPA) and the Brier score, to weight the probability of ancestral nucleotides and, consequently, to determine the most optimal discrete (non-alternative) ancestral states, and 2) the fastest standard and simplified (maximum a posteriori) discreet (non-alternative) MAP approach to reconstruct ancestral states, which reconstructs the ancestral nucleotide just like the nucleotide with the greatest local probability for the alignment site and the tree node^61^. To simplify the comparison of different approaches, we used time-reversible GTR (Tavaré S (1986). Lectures on Mathematics in the Life Sciences. 17: 57–86) and non-reversible RY10.12 model^48^ (the best fitted model to our datasets) in our reconstruction by IQTREE 2.2.0. For the PASTML we used the following parameters: MPPA, the Brier score aware reconstruction based on the HKY model^62^. To be as coherent with the neutrality of the mutations as possible and at the same time to work with most reliable data, we have implemented a search for optimal threshold parameters on both CYTB and ND1 vertebrate data sets. First, we unambiguously showed that only classes 5 and 6 of synonymous site-rate heterogeneity (data categorized into 6 site-rate classes) have similar mutation spectra, but other (more conservative) synonymous site-rate heterogeneity classes have strongly different mutation spectra (Fig. S3). Such a difference in mutation spectra can be driven by negative natural selection. Second, we compared the mutation spectra, computed based on 10 classes of mutation probabilities, and obtained an unambiguous result that mutations with probabilities < 0.3 give the most strongly distinct mutation spectra, regardless of whether all site-rate heterogeneity classes were taken into analysis or only the most varying site-rate classes 5 and 6 were analyzed (Fig. S4). Furthermore, the inclusion of mutations with probabilities < 0.3 introduces a significant variation between our approach and PASTML (Fig. 4; Fig. S2). Therefore, we recommended to filter out all substitutions with probabilities below 0.3, which are usually located on the deep internal branches of the phylogenetic tree, due to inefficiency of deep inner ancestral states reconstruction. Third, we show that the inclusion of all synonymous site-rate heterogeneity classes into the mutation spectra computation significantly reduced the cosine similarities between our and PASTML mutation spectra for synonymous sites as compared to cosine similarities between our and PASTML mutation spectra for all (synonymous and non-synonymous) sites (Fig. S5). Fourth, we additionally demonstrated that more strict exclusion of the data deficient branches (less than 20 mutations per branch and less than 16 from 192 types of substitutions in trinucleotide context) reduced the cosine similarities between our and PASTML mutation spectra for synonymous sites too (Fig. S6). Thus, we strongly recommend users to remove ancestral states with lowest probabilities from all computations (p < 0.3), and also remove all synonymous sites, which have significant evolutionary restrictions on variation (for gamma distribution of site-rate heterogeneity categorized into 6 rate categories, we recommended use rate categories 5 and 6 only) in order to be as consistent as possible with the neutrality of the mutations and simultaneously to work with data with the utmost confidence.

**Fig. 4.**
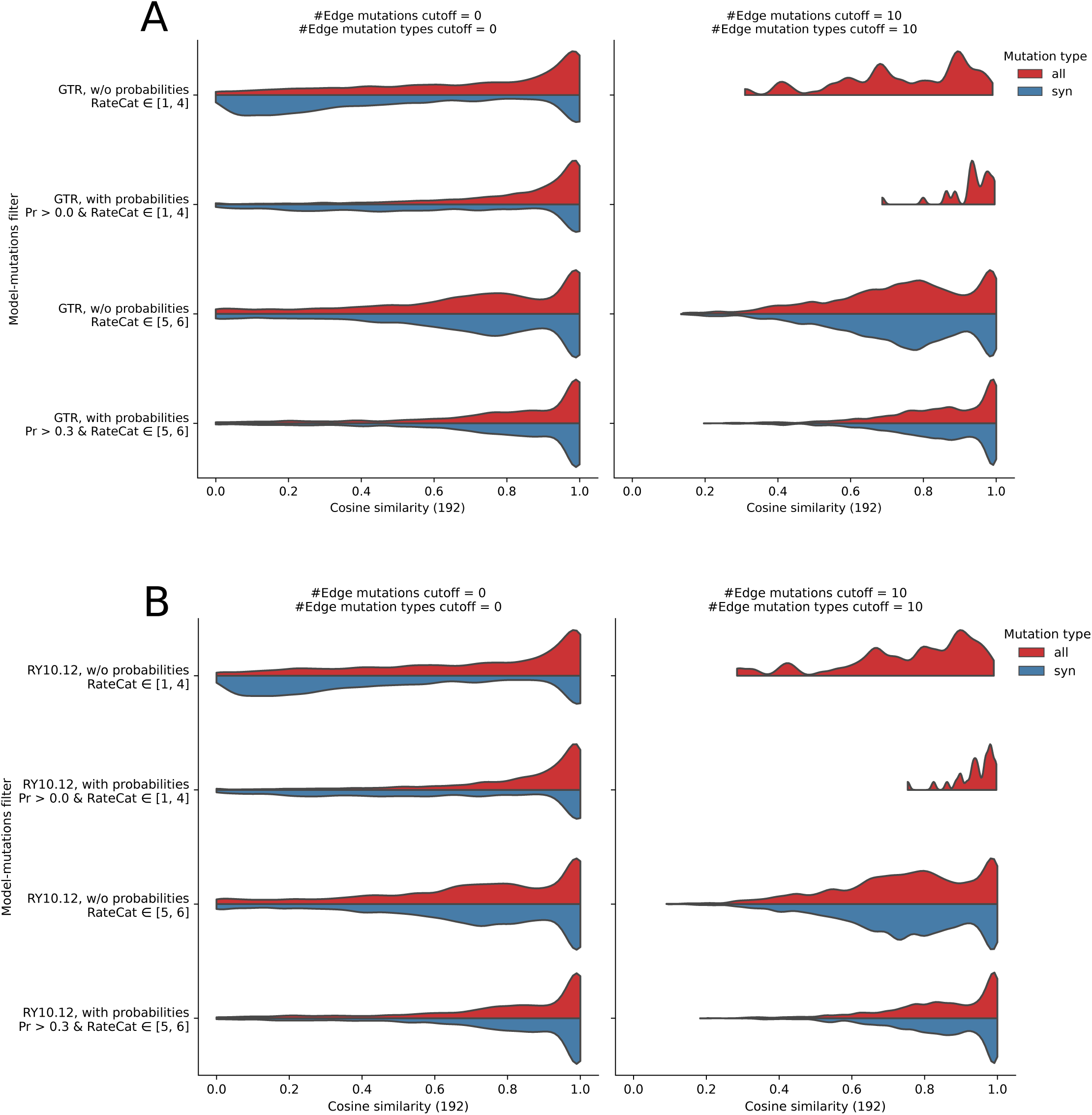
The distributions of pairwise cosine similarities of ND1 Vertebrate mutation spectra computed by PASTML (MPPA, the Brier score aware reconstruction based on HKY model) versus our two approaches: under GTR time reversible model (upper panel) and RY10.12 time non-reversible model (bottom panel). The distributions of pairwise cosine similarities from comparison of mutation spectra for all branches and mutations occurred in alignment sites of two rate categories, [1..4] and [5,6], demonstrated on the 1,2 and 3,4 rows, respectively. Left column shows the distributions of all 192-component mutation spectra, no filters applied. Right column shows the distributions of the 192-component mutation spectra based on tree branches having at least 10 mutations of 10 types. Odd rows demonstrate comparison of our simple ancestral states inference (MAP reconstruction of single/best ancestral state). Even rows demonstrate comparison of mutation spectra considering probabilities of all possible ancestral states.

In general, we demonstrated the effectiveness of our “uncertainty principle” normalization approach with the defined thresholds (see above) applied, we showed that this fast computational approach (∼1000 times faster than PASTML) provides similar mutation spectra result as compared to one reconstructed by the best but slowest general ancestral approach to date, PASTML.

Not only the computation speed but also the accuracy and stability of computations are crucial for obtaining precise mutation spectra, hence it is important to pay critical attention to the convergence between the observed and expected mutation spectra obtained through evolutionary simulations or, in other words, the convergence between the spectra of simulated and reconstructed ancestral states. In order to test this, we performed several types of evolutionary simulations using IQTREE 2.2.2.7 AliSim tool^63^. We took two modes of evolutionary simulations: 1) based on the time-reversible GTR model of nucleotide substitution and lognormal distribution (μ=-2, σ=1) of model parameters, and 2) based on non-reversible 12.12^48^ model of nucleotide substitution and normal distribution (mean=0, stddev=0.5) of model parameters (for these purposes we used standard random python library). We used either the natural sequence of the human CYTB gene or the random nucleotide sequence of equal length (1140 b.p.) as the root sequence of each simulation. For all simulations we used gamma distribution of site-variation rates categorized into 6 rate categories (option ‘+G6’). The empirical base frequencies (option ‘+F’) were taken into consideration in every simulation, the base frequencies were sampled as 4 random values (by random python library) and normalized by unity sum rule. We also randomized the fraction of invariable sites in alignment (option ‘+I’) from 0 to 0.5. We generated two types of random birth-death trees: shallow (100 leafs per tree) and deep (1000 leafs per tree). To normalize branch tree lengths we used the ‘rlen’ option with min=0, median=0.001 and max=0.01. In order to simulate outgroup for each tree we computed the mean length from root to leaf and multiplied it by 5. For each combination of model, size of tree and root sequence type we generated 100 replicas, for a total of 800 replicas. The results of simulations demonstrated on Fig 5, the convergence between the simulated (ground truth) and reconstructed mutational spectra (ancestral states) are rather precise, even considering the extreme modes of nucleotide evolution. This indicates that it is possible to precisely analyze almost any complex case of species-specific DNA evolution using the approach we have chosen.

**Fig. 5.**
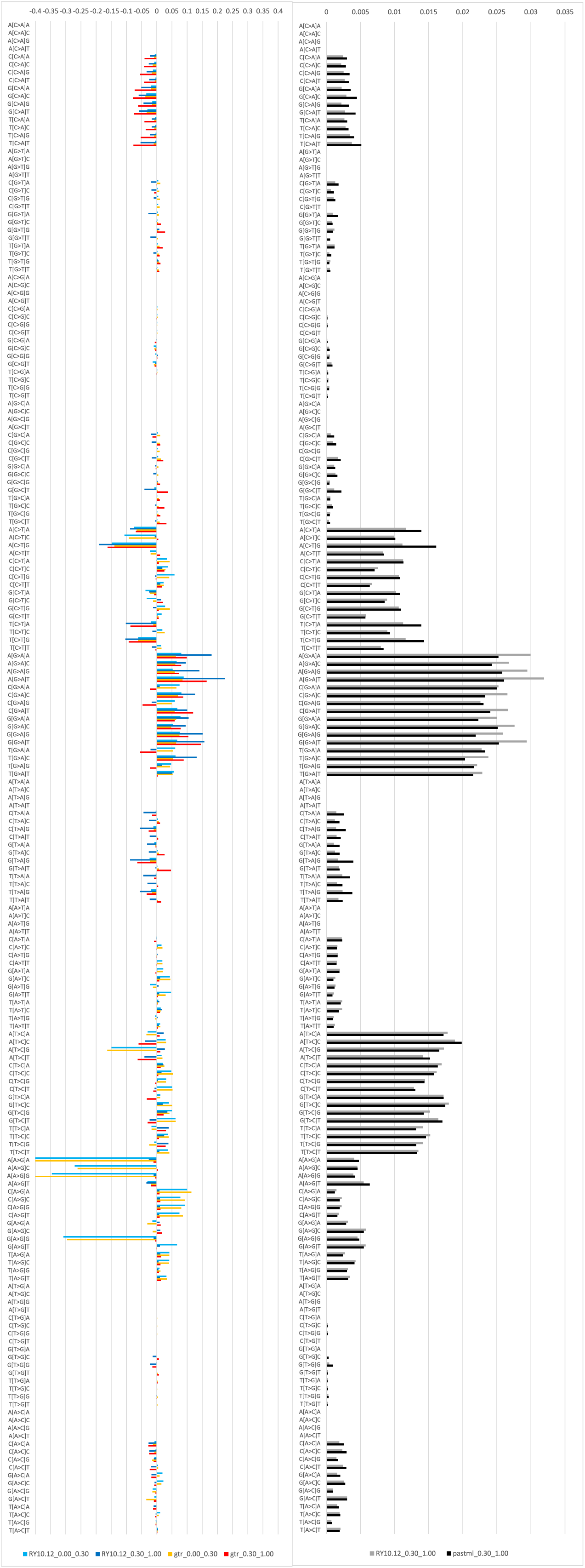
Testing for ancestral states and/or mutational spectra computational accuracy on random trees: the comparison between simulated and reconstructed 192 component. (A) and 12 component (B) mutational spectra based on cosine similarities.

The uniform distribution of branch lengths prevents the generation of random birth-death trees from being a case of real evolution, purely speaking. So, to test the accuracy of our mutational spectra inference approach, we use a more rigorous example: our second simulations was based on the actual tree topology of 4705 mammalian species [timetrees/01_step2/4705sp_mammal-time.tree file from^64^]. In this type of simulation we also used either the natural sequence of the human CYTB gene or a random sequence (that is identical length as CYTB (1140 b.p.)) as the root sequence for each simulation. Random sequences, approximated in nucleotide composition to the natural sequences, were generated considering Shannon entropy. Only sequences whose entropy differed from the entropy of human CYTB frequencies by no more than 0.05 were used. (Fig. S7). We also took two modes of evolutionary simulations, the time-reversible GTR model and the non-reversible 12.12 model, as described above. In these simulations, in order to be as precise as possible, we 1) change the cosine similarities by Euclidean distances for the comparison between the simulated (ground truth) and reconstructed mutational spectra, 2) analyze this distances for each of the alignment site variability class separately, and 3) analyze synonymous mutations and all mutations separately. We use the variability class prediction of each alignment site based on gamma rate heterogeneity across alignment sites by IQTREE 2.2.2.7^53,54^. Cosine similarity only takes into account the angle between vectors, while Euclidean distance takes into account their weight or magnitude. In other words, we consider the exact difference in mutation numbers on each tree branch between ground truth and reconstructed data in these simulations. Despite the undoubted importance of Euclidean distances for mutation spectra reconstruction tests, they are not standardized as cosine similarities. To standardize Euclidean distances, or in other words, to obtain neutral expectation of Euclidean distances, we compute Euclidean pairwise distances between mutational spectra in simulated (ground truth) replicas. The results of reconstructions (observations) and standardized Euclidean distance distributions for synonymous mutations only demonstrated on Fig 6 (data for all mutations, synonymous and non-synonymous, shown in Fig. S8). It is very important that 1) the Euclidean distance distributions are multimodal for every alignment site variability class in 192 component mutational spectra, 2) only the most variable site class (cat6) have the largest and leftmost intersection between neutrally expected and observed Euclidean distance distributions, and 3) compared to a neutral expectation, the Euclidean distances of the majority of less variable classes of alignment sites are significantly biased towards zero (Supplementary Files 2 and 3). These facts prove that only a portion of most variable synonymous substitutions can be classified as neutrally evolved cases, as expected in true natural conditions.

**Fig. 6.**
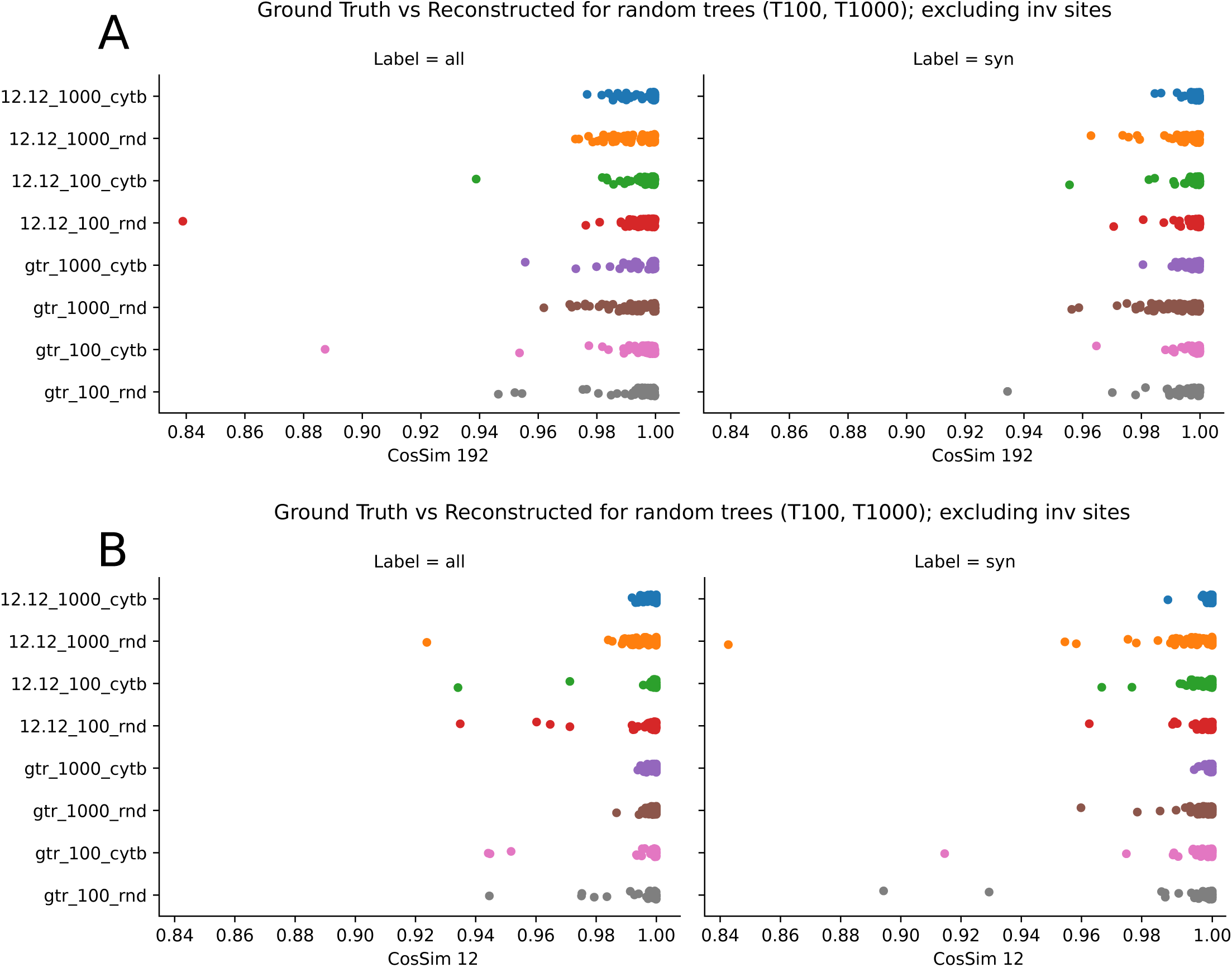
Testing for ancestral states and/or mutational spectra computational accuracy on a real tree of 4705 mammalian species: the comparison between simulated and reconstructed 192 component (bottom row) and 12 component (top row) mutational spectra calculated using only synonymous mutations based on Euclidean distances.

### (5) Accurate mutational spectra inference from reconstructed ancestral states probabilities: an application for inter-species mitochondrial gene data

After carefully reconstructing and filtering ancestral state probabilities in each alignment site of each inner tree node, we infer the mutational spectrum. In cancer studies to build mutation spectra, it is common to normalize the frequencies of observed mutations using the trinucleotide frequency in the human exome or genome^9^ or using the trinucleotide frequency of genes under study. On a comparative-species scale differences in nucleotide content of species could significantly affect the rate of observed mutations (for example in AT rich genome many mutations would be from A and T) and thus should be taken into account carefully. To do this, we normalize the frequencies of the mutations observed in the trinucleotide content (taking into consideration ancestral states probabilities) using two alternative schemes: 1) using the frequency of trinucleotides (trinucleotides centered on synonymous sites) of all sequences (ancestral and existing) of the phylogeny analyzed, we do so for compatibility with cancer data stored in the COSMIC database^65^, 2) using the frequency of trinucleotides (trinucleotides centered on synonymous sites excluding conservative ones) from all possible synonymous alternatives of all sequences (ancestral and existing) of the phylogeny analyzed, we make it on the basis of the simple evolutionary logic of Nei and Gojobori^66^. Comparing these two normalization procedures demonstrates that the results obtained are identical on our datasets (Fig. S9).

The most intriguing question is how does our approach work with respect to specific types of substitutions, is it able to differentiate between less biologically likely substitutions and more biologically likely substitutions? To this end, we chose the most challenging case, the evolution of mitochondrial DNA on an inter-species scale, which is known to be qualitatively non-equal on various strands. Heavy (H) mtDNA strand (as compared with light (L) mtDNA strand) is known to have encountered elevated mutagenesis (deamination of cytosine and adenine, oxidation of guanine, etc) due to its existence in single-strand mode over a significant period of time. Because of this mutation imbalance on the different mtDNA strands, complementary substitutions should have an unequal frequency (in the top 3 mutation pairs, in L mtDNA strand notation, e.g., G→A >> C→T, T→C >> A→G, C→A >> G→T), skewed by frequent nucleotide damage on the H strand^67–69^. This mutational feature has been demonstrated to require non-reversible Markov models for analysis^67^. In Fig. 7A we show that in comparison with PASTML (only time-reversible Markov models available – MPPA, the Brier score aware reconstruction based on HKY model), our approach (both GTR time reversible Markov model and RY10.12^48^, the best fitted non-reversing Markov model) makes it possible to (Fig. 7A): 1) maximize inequality between frequencies of complementary substitutions of major mutation types (G→A >> C→T, T→C >> A→G^67–69^), 2) correctly differentiate the unequal frequencies of complementary substitutions of minor mutation types (C→A >> G→T, G→C > C→G, A→C >> T→G^68^), and 3) correctly estimate the relative low frequency of minor mutation types as compared to major mutation types (for example, G→C & C→G in comparison with C→T & G→A^68^). This allows us to reconstruct the mutation spectrum more precisely and therefore more qualitatively and quantitatively biologically appropriate. Additionally, the comparison of mutation spectra obtained on both datasets, ND1 and CYTB, (Fig 7 and Fig. S10, respectively) shows that best fitted non-reversible Markov model and GTR reversible Markov model for ancestral synonymous mutations reconstruction give us qualitatively similar estimates of mutation frequencies in trinucleotide context. First, this indicates the robustness of our mutational spectra estimates. Second, it shows that if only qualitative estimation of the mutational spectrum is desirable to the user, it is possible to use a time reversible Markov model of nucleotide substitution (e.g. GTR) rather than a non-reversible model.

**Fig. 7.**
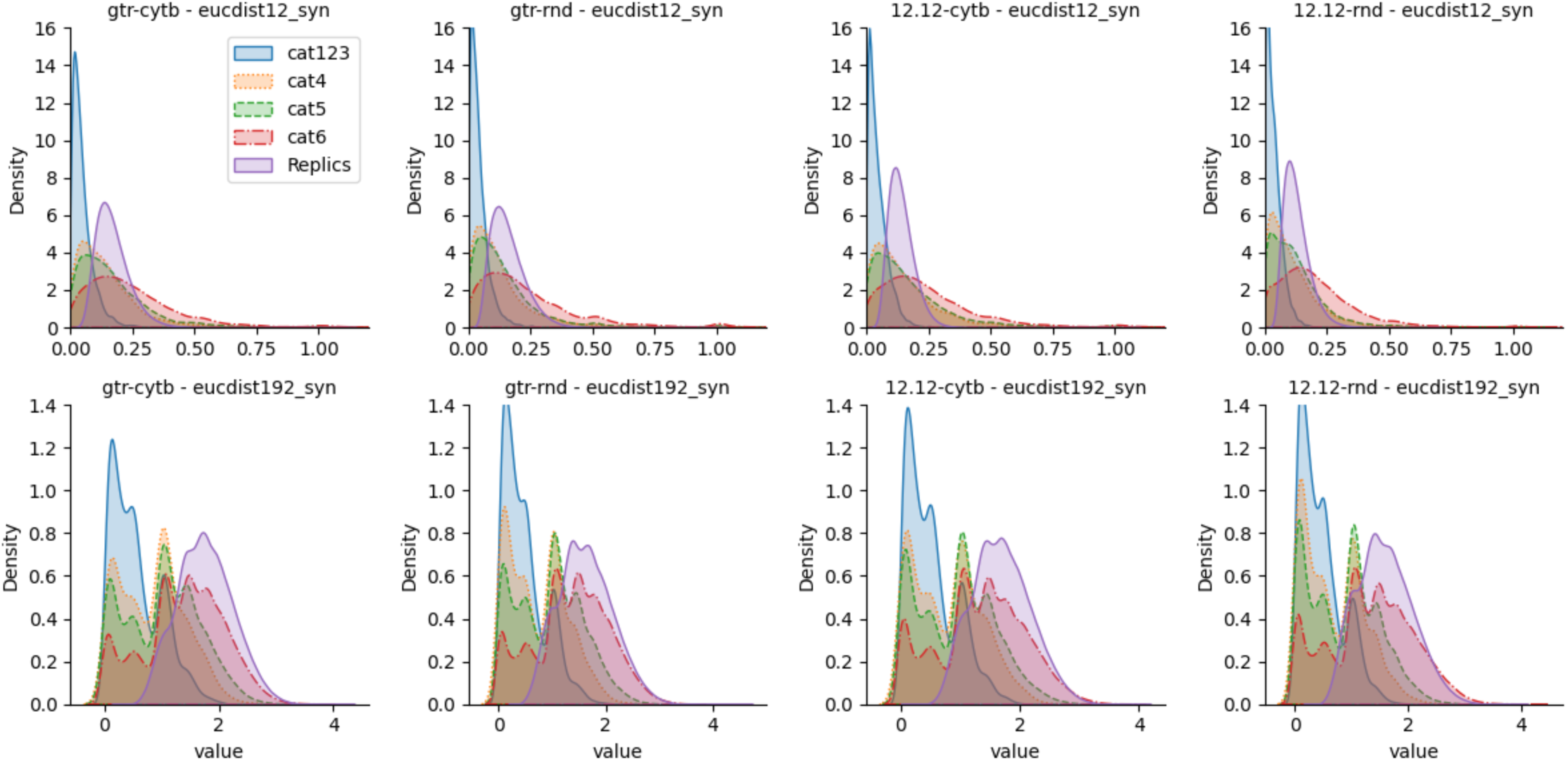
The Mammals ND1 mutation spectrum, RateCat [5, 6] used. Left panel, the comparison of spectra obtained from PASTML-reconstructed ancestors with our approach: red and orange bars correspond to GTR substitution model (red, p>0.3; orange, p<0.3), blue and cyan bars correspond to RY10.12 substitution model (blue, p>0.3; cyan, p<0.3). Right panel, the Mammals ND1 mutation spectra obtained by PASTML (black) and by our approach, RY10.12 substitution model (gray); both our and PASTML reconstructions based on p>0.3 ancestral probability class.

### (6) Comparison between observed nearly neutral mutational spectrum and expected fully neutral mutational spectrum expectations

Currently, there is a broad range of codon evolution Markov models known as MutSel models that specifically separate the effects of neutral mutagenesis from selection based on the Fisher-Wright mutation selection framework^28,70–78^. In this respect, we implemented an additional custom pipeline feature, we can compare the observed nearly neutral mutational spectrum with the fully neutral predicted spectra. By doing so, we are able to identify and subtract even the smallest effects of natural selection. We implemented this step based on the most powerful type of MutSel Markov model of codon evolution that made it possible to have a selective coefficient even on each synonymous change^79^. This MutSel model implemented in Pyvolve package^80^ that supports site-wise heterogeneity (including setting of invariable sites), phylogenetic branch lengths heterogeneity, the specification of a custom rate matrix for simulation, and custom set of states to evolve. We upgraded the pyvolve package to add support for different genetic codes. Package version used in the simulations is available at https://github.com/kpotoh/pyvolve. Thus, based on pyvolve it is simple to specify arbitrary models, and programmatically evaluates the effect of selection for each type of nucleotide substitutions in the observed mutation spectrum. Thanks to the maximal number of available settings in pyvolve, it is easy to simulate the neutral evolution of natural sequences on the natural phylogenetic tree, which enables a direct comparison of the observed and expected mutational spectrum (e.g., by subtracting the anticipated expected mutational spectrum from that observed, we can estimate the proportion of the mutational spectrum associated with even tiny selection). This, finally, could significantly improve the result and simplify the biological interpretation of mutation spectrums as well. For example, in Fig. 8 we compared 200 simulated neutral mutational spectra of the CYTB mouse gene with observed ones, it is clearly demonstrated that various substitutions in multiple DNA motifs are statistically under-represented or over-represented in the observed spectrum relative to expected neutral spectra. This in turn allows us to conclude that the comparison of the observed mutational spectra with simulated neutral spectra based on the MutSel model can provide evidence for even tiny selection pressure on specific mutated DNA motifs/trinucleotides.

**Fig. 8.**
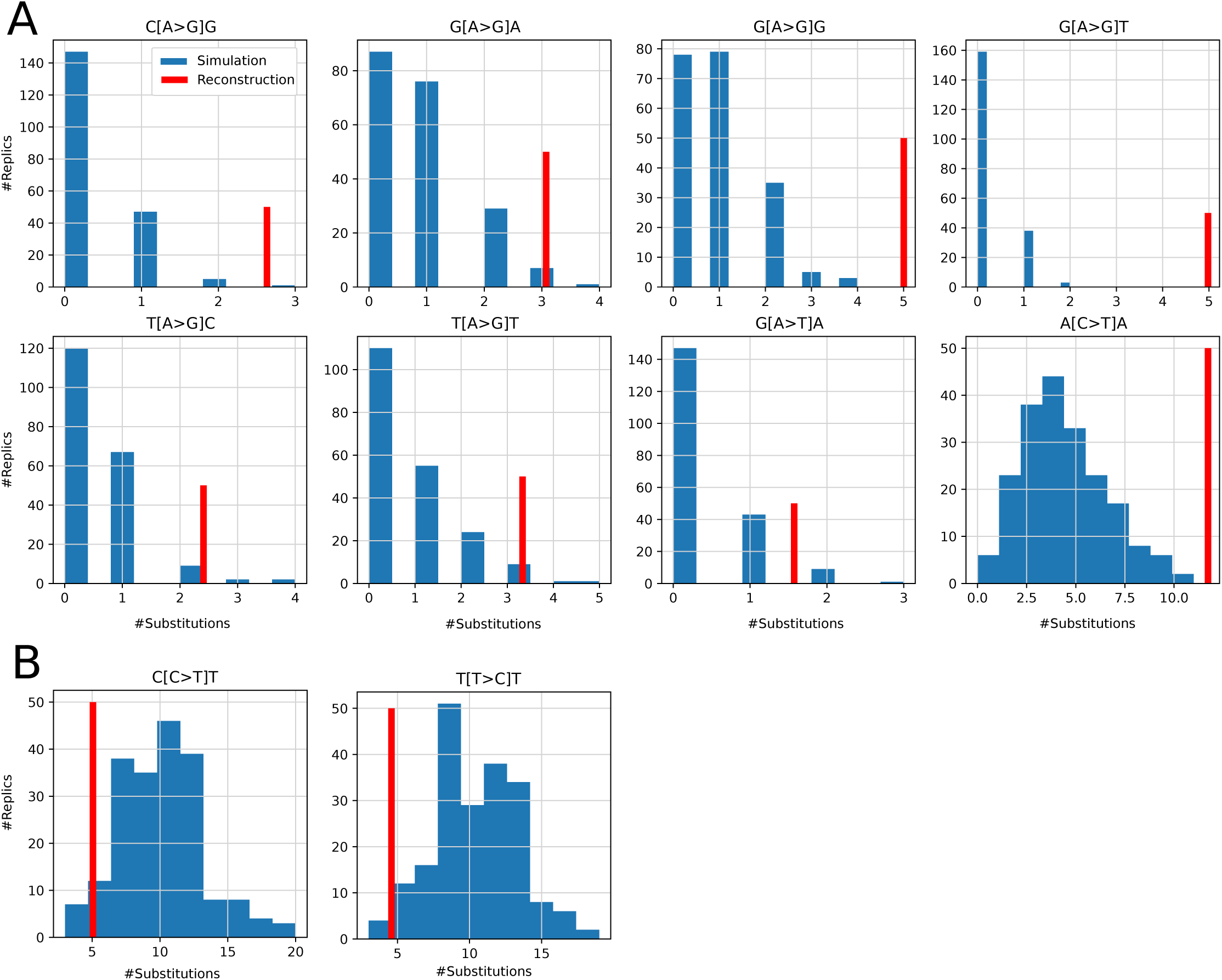
The comparison of 200 simulated neutral mutational spectra of the CYTB mouse gene with observed ones. (A) DNA motifs and substitutions statistically over-represented in the observed spectrum relative to expected neutral spectra (p<0.05); (B) DNA motifs and substitutions statistically under-represented in the observed spectrum relative to expected neutral spectra (p<0.01).

## CONCLUSION

Thus, we have made NeMu a flexible, convenient and interactive pipeline that makes it possible to accurately disentangle neutral mutation spectra from evolutionary data. Its flexibility makes it possible to estimate mutation spectra specific to individual species and higher taxa. Since the neutral mutation spectrum is the focus of most evolutionary studies, the NeMu pipeline could be the software that biologists need today.

## DATA AVAILABILITY

Repository https://github.com/mitoclub/nemu-pipeline contains pipeline, analyses code and pipeline documentation.

## NEMU PIPELINE ACCESS

https://biopipelines.kantiana.ru/nemu/

## AUTHOR CONTRIBUTIONS

The design of the study developed by K.G., the pipeline implemented by B.E. and K.G. Data mining, processing and analysis performed by B.E. and K.G. Web-interface developed by B.E. Manuscript prepared by K.G, B.E. and K.P. All authors (B.E., K.P. and K.G.) discussed in depth the manuscript and the rationale behind the project. All authors read and approved the final manuscript.

Conflict of interest statement. None declared.

