## Appendix and Supplementary figures for "NeMu: A Comprehensive Pipeline for Accurate Reconstruction of Neutral Mutation Spectra from Evolutionary Data"

**Supplementary Materials to the manuscript of Efimenko et al. "NeMu: A Comprehensive Pipeline for Accurate Reconstruction of Neutral Mutation Spectra from Evolutionary Data"**

APPENDIX 1

For example, the evolution of DNA can be simply mathematically formalized as a system of ordinary differential equations showing changes of all nucleotide frequencies (A, G, C and T) in time; here in a time-reversible manner:

$$\begin{cases} f_A{}'(t) = -f_A(t)\mu_A + \sum\limits_{x \neq A} f_x(t)\mu_{Ax}dt, \\ f_G{}'(t) = -f_G(t)\mu_G + \sum\limits_{x \neq G} f_x(t)\mu_{Gx}dt, \\ f_C{}'(t) = -f_C(t)\mu_C + \sum\limits_{x \neq C} f_x(t)\mu_{Cx}dt, \\ f_T{}'(t) = -f_T(t)\mu_T + \sum\limits_{x \neq T} f_x(t)\mu_{Tx}dt, \end{cases}$$

where $\mu_{yx} = \mu_{xy} \ \ (y \neq x)$

For example, in matrix form it could be rewritten as:

$$Q = \begin{bmatrix} -\mu_A & \mu_{AG} & \mu_{AC} & \mu_{AT} \\ & -\mu_G & \mu_{GC} & \mu_{GT} \\ & & -\mu_C & \mu_{CT} \\ & & & -\mu_T \end{bmatrix} = \begin{bmatrix} -0.049 & 0.04 & 0.006 & 0.003 \\ & -0.049 & 0.006 & 0.003 \\ & & -0.072 & 0.06 \\ & & & -0.072 \end{bmatrix}.$$

We can use Wolfram Mathematica software to simple solve this system of ordinary differential equations:

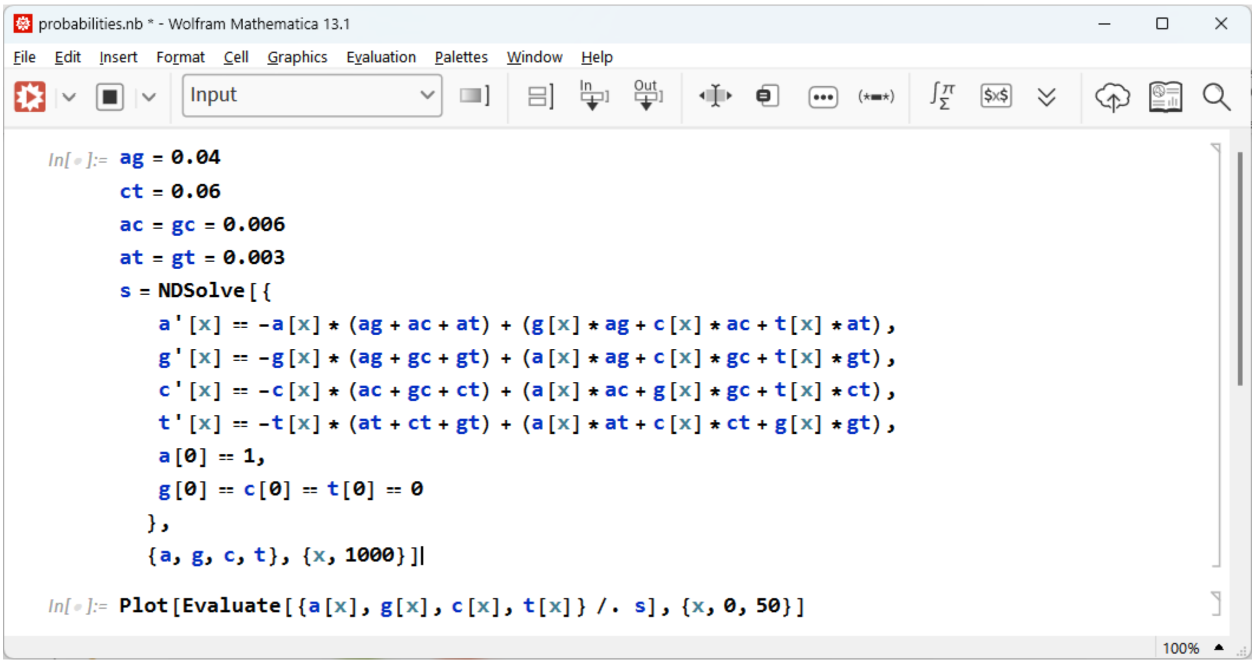

```
In[ ]:= ag = 0.04
        ct = 0.06
        ac = gc = 0.006
        at = gt = 0.003
        s = NDSolve[{
           a'[x] == -a[x] * (ag + ac + at) + (g[x] * ag + c[x] * ac + t[x] * at),
           g'[x] == -g[x] * (ag + gc + gt) + (a[x] * ag + c[x] * gc + t[x] * gt),
           c'[x] == -c[x] * (ac + gc + ct) + (a[x] * ac + g[x] * gc + t[x] * ct),
           t'[x] == -t[x] * (at + ct + gt) + (a[x] * at + c[x] * ct + g[x] * gt),
           a[0] == 1,
           g[0] == c[0] == t[0] == 0
          },
          {a, g, c, t}, {x, 1000}]

In[ ]:= Plot[Evaluate[{a[x], g[x], c[x], t[x]} /. s], {x, 0, 50}]
```

As a result, we can clearly demonstrate that the more the time of evolution we analyzed, the less the expected differences in nucleotide frequencies we have:

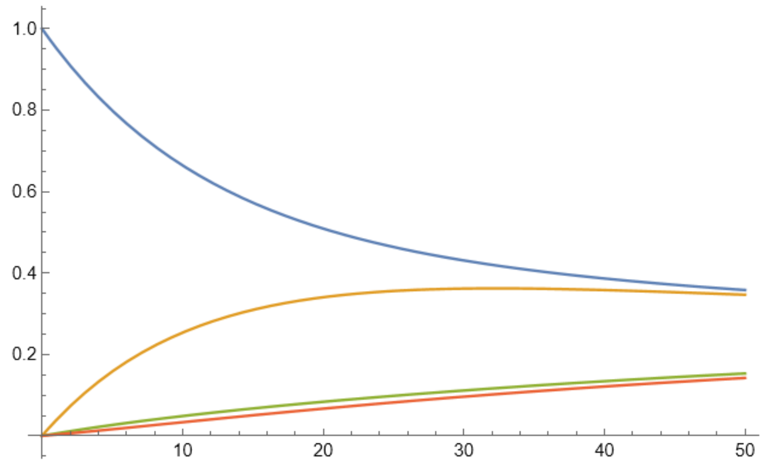


Time-related ancestral nucleotide probabilities according to the substitution model that showed above; from contemporary time (extant nucleotide is A) into old ages:
A – blue;
G – orange,
C – green,
T – red colors.

APPENDIX 2

Scheme of blurring probabilities for ancestral sequence states (it may also represent an imprecision of sequence alignment near the root (doi: 10.1093/molbev/msy055)), from contemporary time (extant nucleotide is A) into old ages (Nx - ancestors).

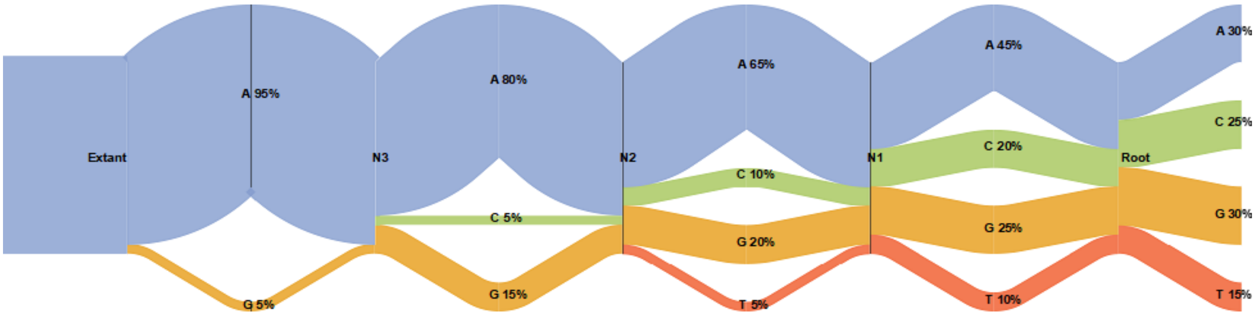

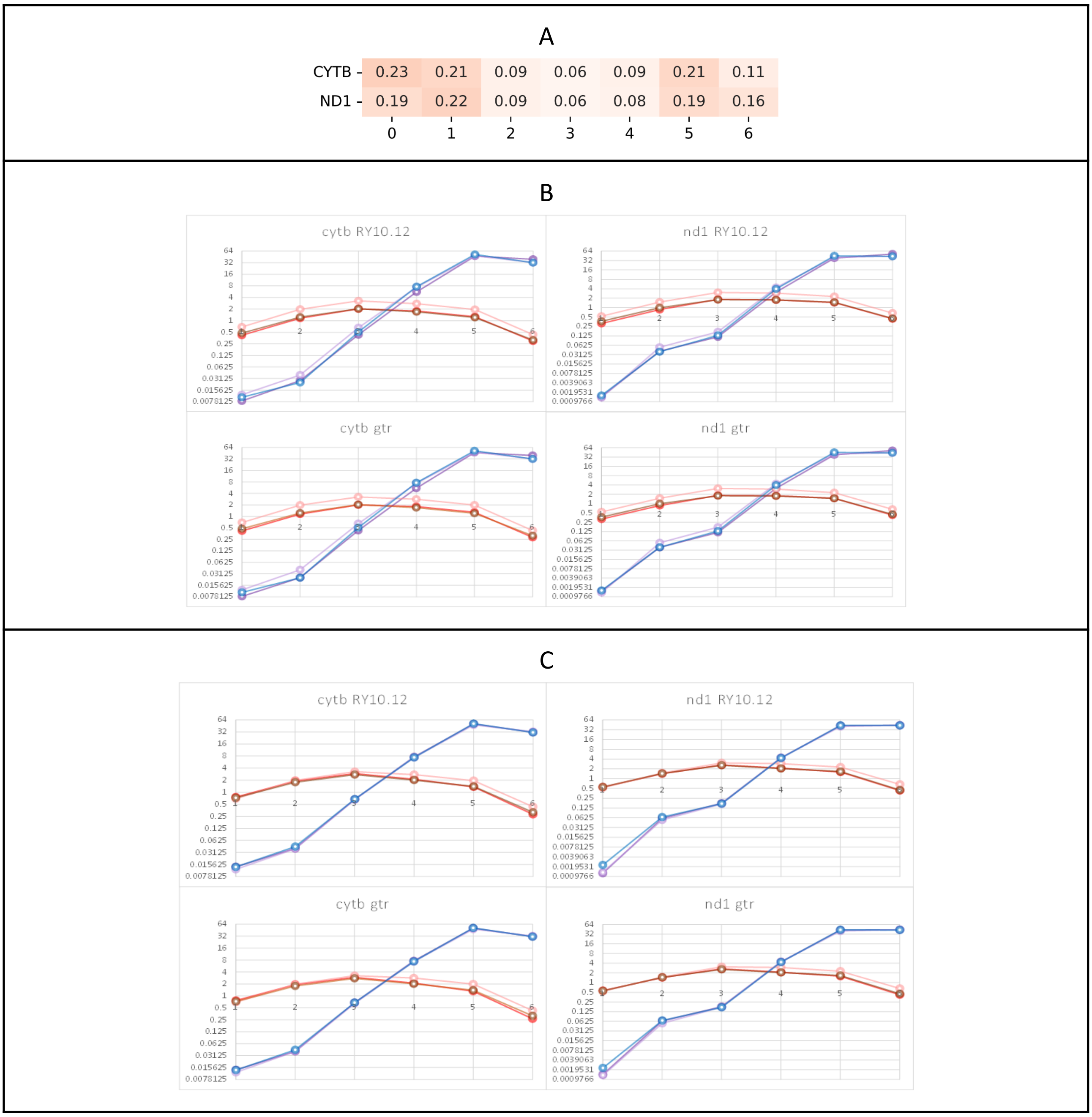


Fig. S1. Variability of alignment sites in mammalian cytb and nd1 mitochondrial genes. (A) The fraction of alignment sites characterizing by 0 - 6 rate heterogeneity categories. Six rate categories (from 0, totally conservative, to 6, totally variable) of gamma distribution were used. (B) The percent (Y-axes) of substitution types and rate heterogeneity classes (X-axes) across alignment sites, all probabilities of mutations. Two substitution types were shown by different colors: synonymous substitutions by purple/blue, and non-synonymous substitutions by red/brown. Saturated and light red and purple color tones show "uncertainty principle" normalization of ancestral states (considering and weighting probabilities of all possible ancestral states) and simple ancestral states inference (MAP reconstruction of single/best ancestral state), respectively. Blue and brown colors show PASTML results (MPPA, the Brier score aware reconstruction based on HKY model). (C) The percent (Y-axes) of substitution types and rate heterogeneity classes (X-axes) across alignment sites, probabilities of mutations > 0.3. Set of colors like in (B).

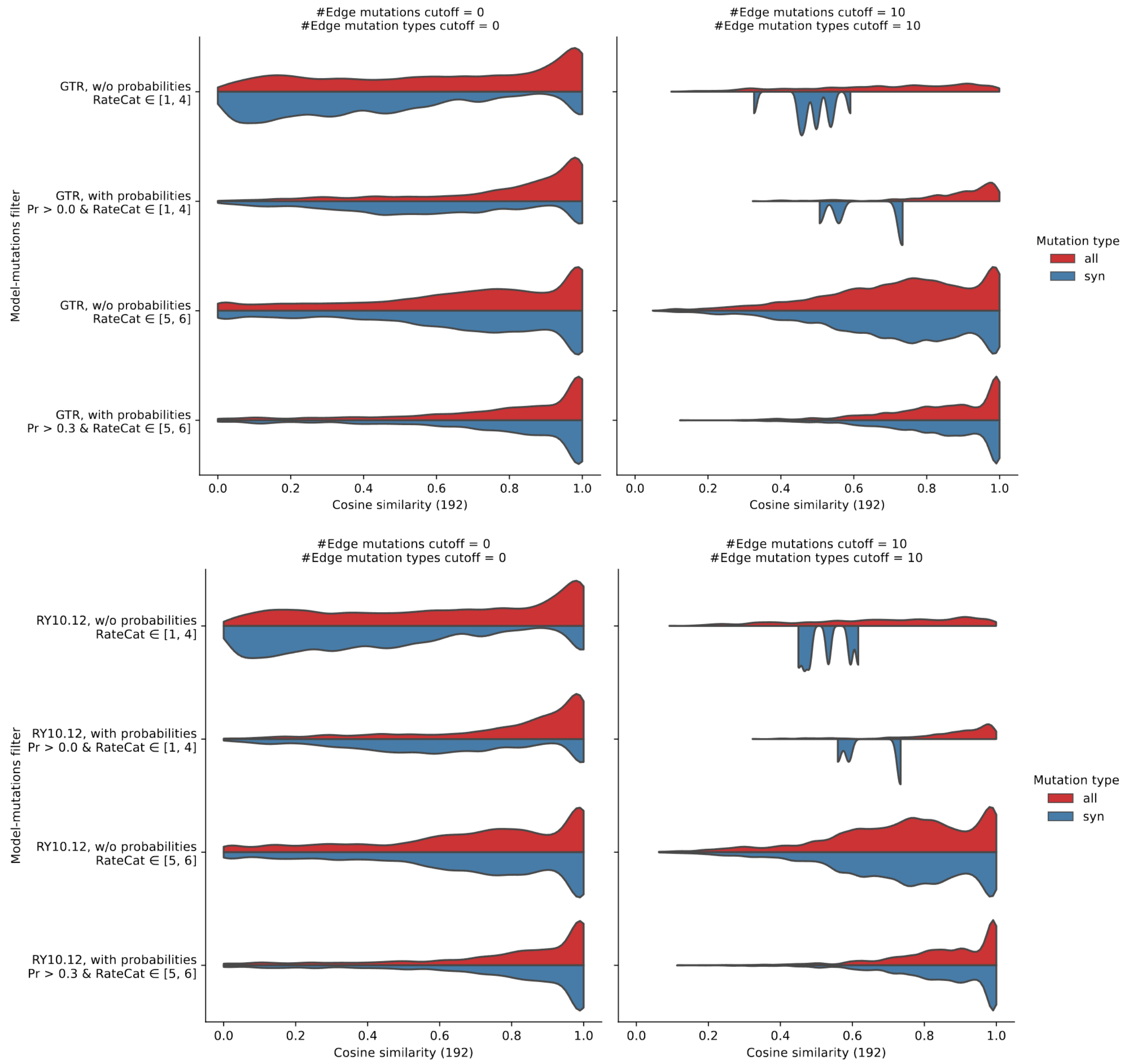


Fig. S2. The distributions of pairwise cosine similarities of CYTB Vertebrate mutation spectra computed by PASTML (MPPA, the Brier score aware reconstruction based on HKY model) versus our two approaches: under GTR time reversible model (top panel) and RY10.12 time non-reversible model (bottom panel). The distributions of pairwise cosine similarities from comparison of mutation spectra for all branches and mutations occurred in alignment sites of two rate categories, [1..4] and [5,6], demonstrated on the 1,2 and 3,4 rows, respectively. Left column shows the distributions of all 192-component mutation spectra, no filters applied. Right column shows the distributions of the 192-component mutation spectra based on tree branches having at least 10 mutations of 10 types. Odd rows demonstrate comparison of our simple ancestral states inference (MAP reconstruction of single/best ancestral state). Even rows demonstrate comparison of mutation spectra considering probabilities of all possible ancestral states.

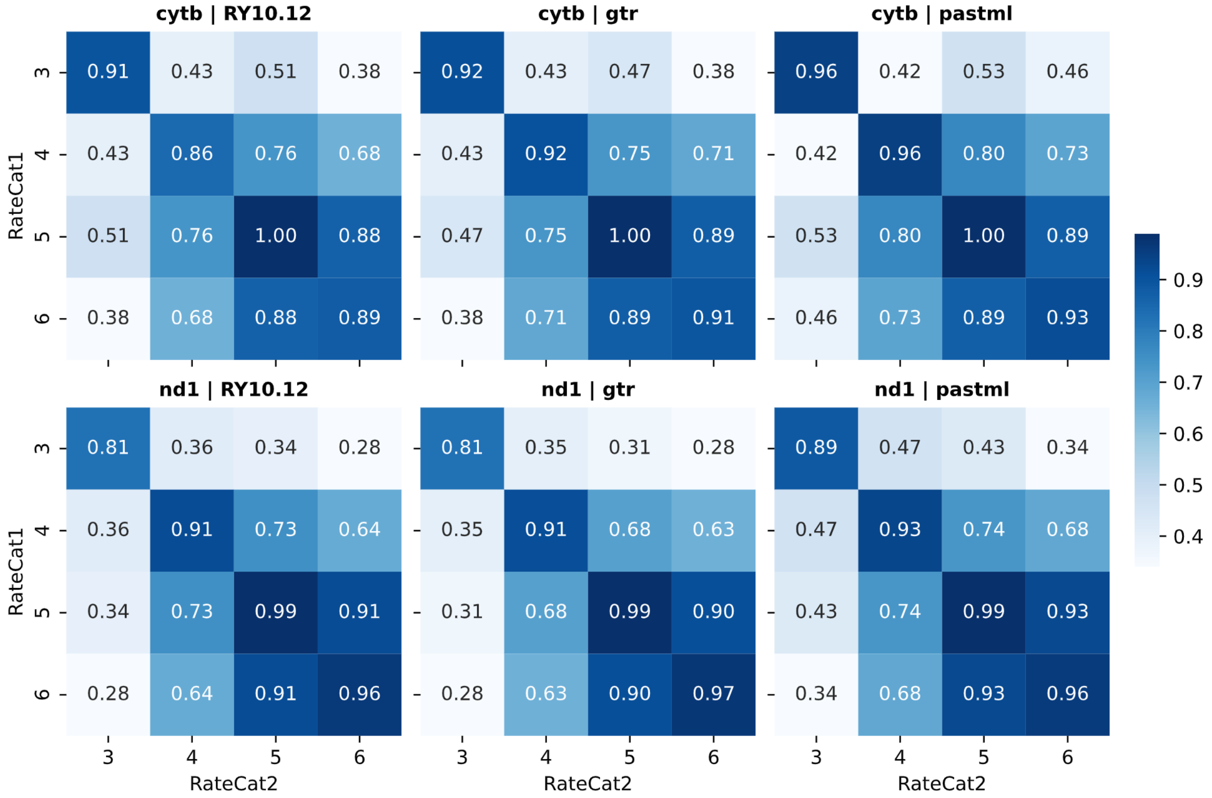


Fig. S3. Pairwise comparisons of mutation spectra (by cosine similarities) computed on the basis of the last four classes of site-rate heterogeneity (invariable class 0 and conservative classes 1 and 2 were omitted due to a lack of data). The median cosine similarities obtained by 50% jackknife were shown.

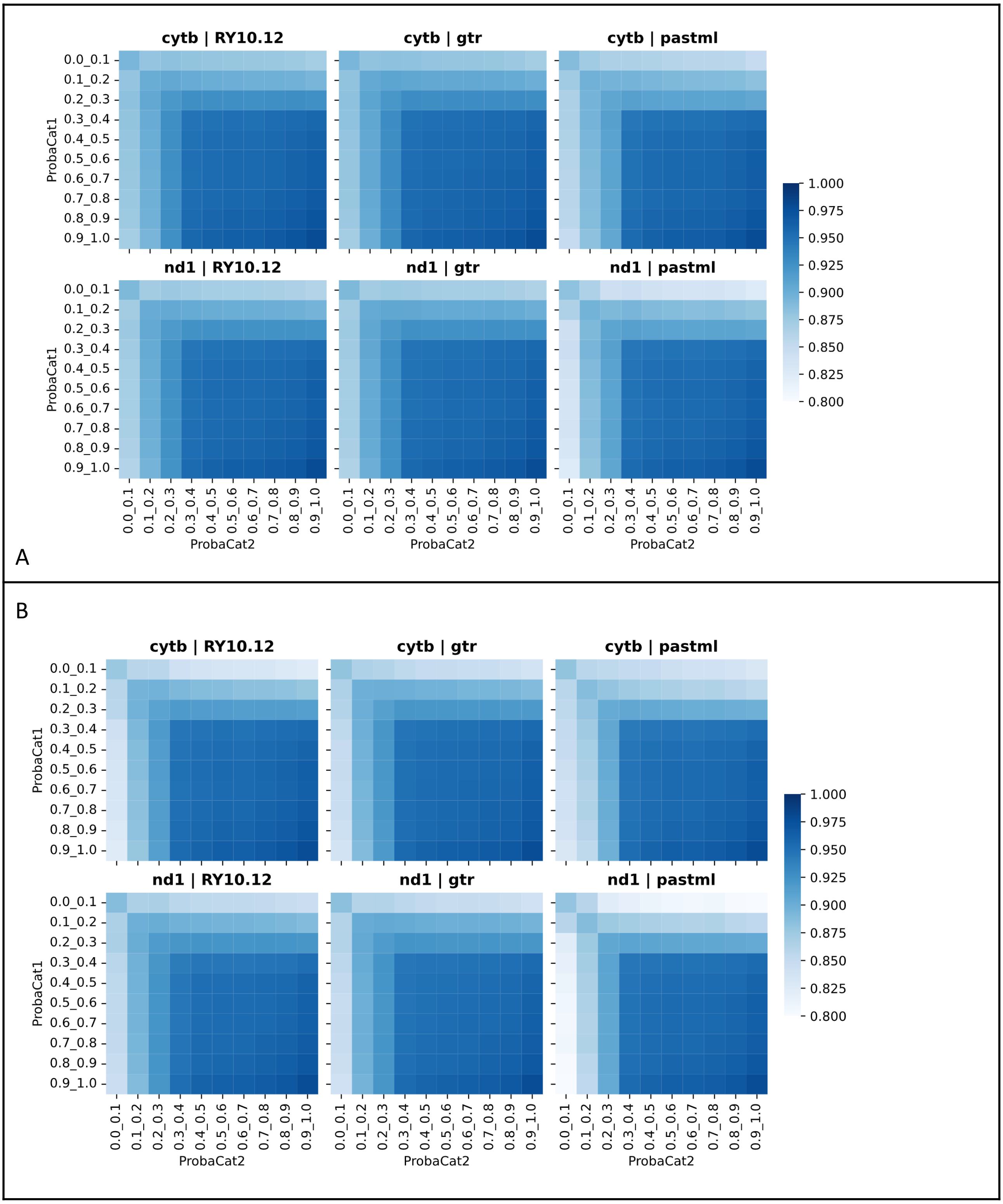


Fig. S4. Pairwise comparisons of mutation spectra (by cosine similarities) computed on the basis of the 10 classes of mutation probability classes. (A) Computation based on all site-rate heterogeneity classes, (B) Computation based on the 5 and 6 site-rate heterogeneity classes. The median cosine similarities obtained by bootstrap were shown. The median cosine similarities obtained by 50% jackknife were colorized.

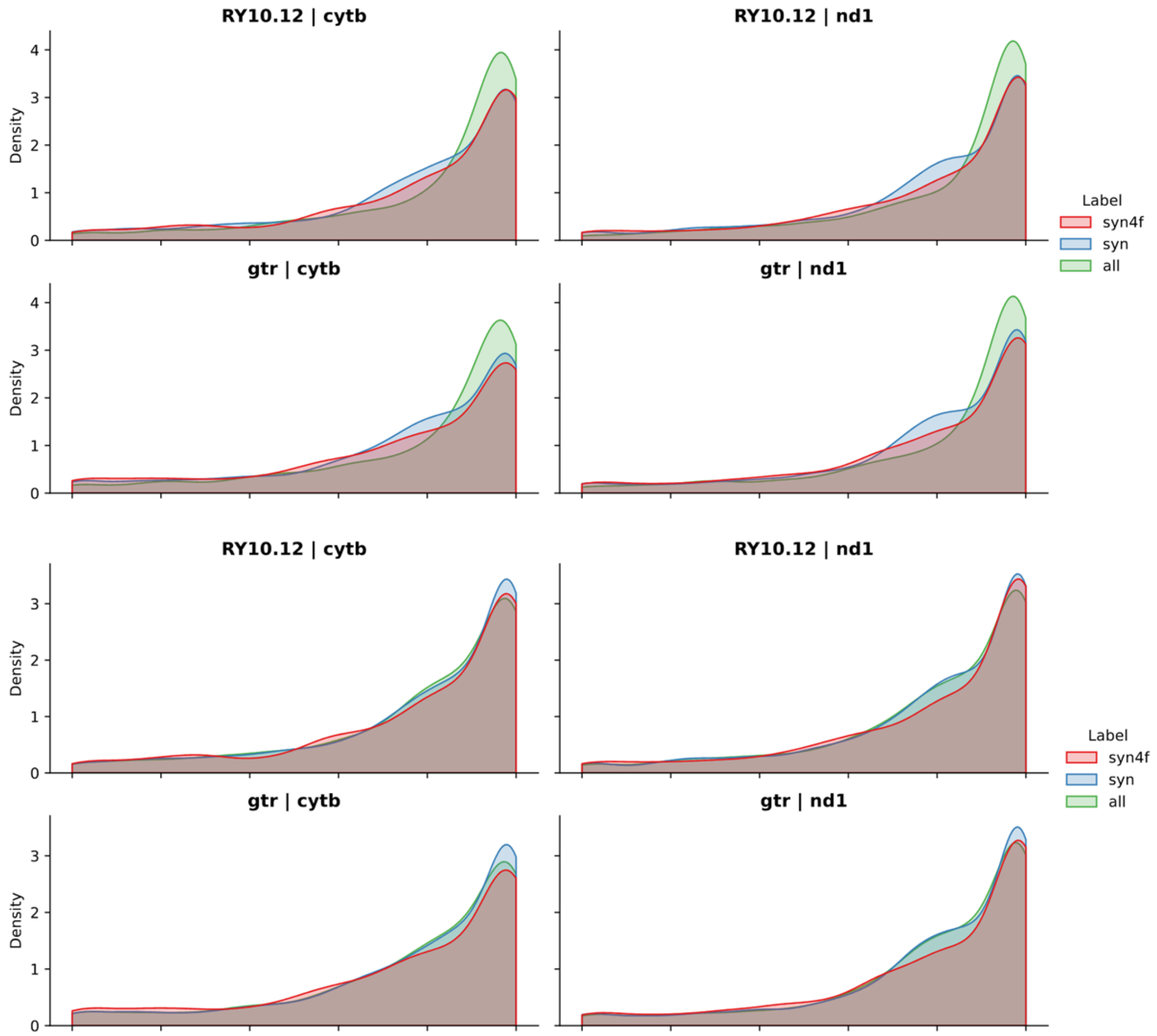


Fig. S5. The effect of inclusion of all synonymous site-rate heterogeneity classes into the mutation spectra computation on cosine similarities between our and PASTML mutation spectra. Upper 4 plots, all site-rate heterogeneity categories under consideration. Lower 4 plots, only 5 and 6 site-rate heterogeneity classes analyzed.

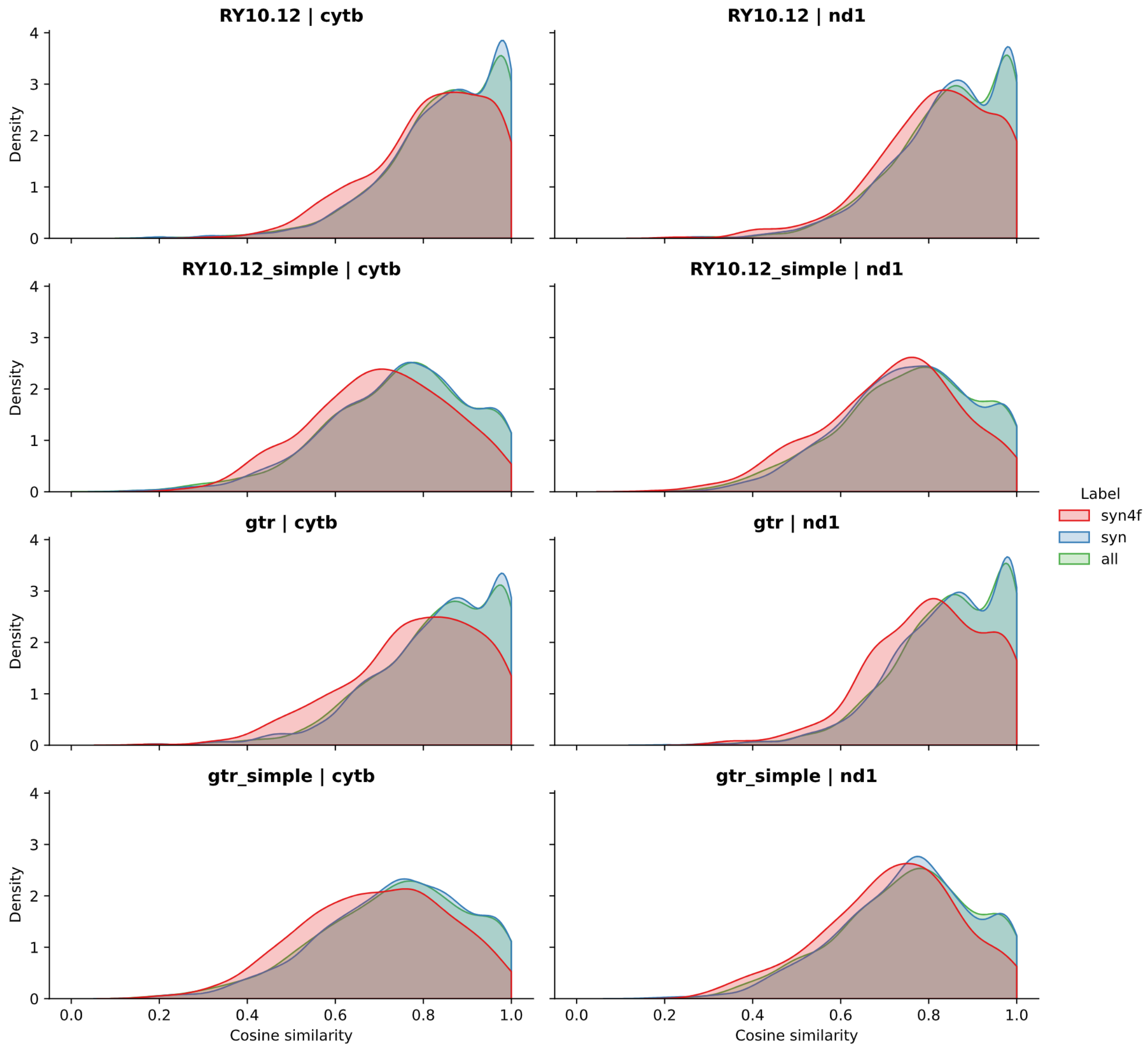


Fig. S6. The distributions of pairwise cosine similarities of CYTB (left column) and ND1 (right column) Vertebrate mutation spectra (as in Fig. 4, FigS2), the probabilities of mutations >0.3, only 5 and 6 site-rate heterogeneity classes, short branches with data deficiency (less than 20 mutations per branch and less than 16 from 192 types of substitutions in trinucleotide context) have been omitted.

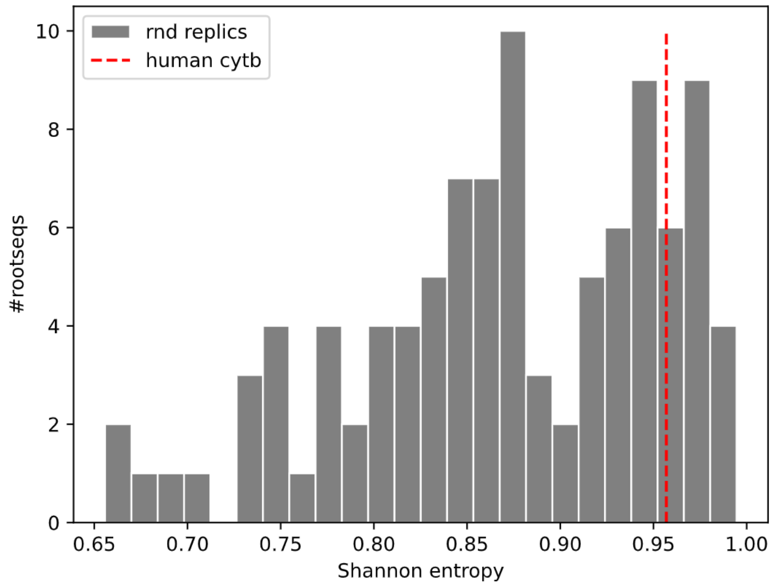


Fig. S7. The distributions of Shannon entropies calculated for random root sequences in 100 replicas based on nucleotides frequencies. Red dashed line indicates entropy for the human cytb sequence. Low entropy values denote prevalence of a single or couple of nucleotides in the sequence, i.e. not a natural scenario.

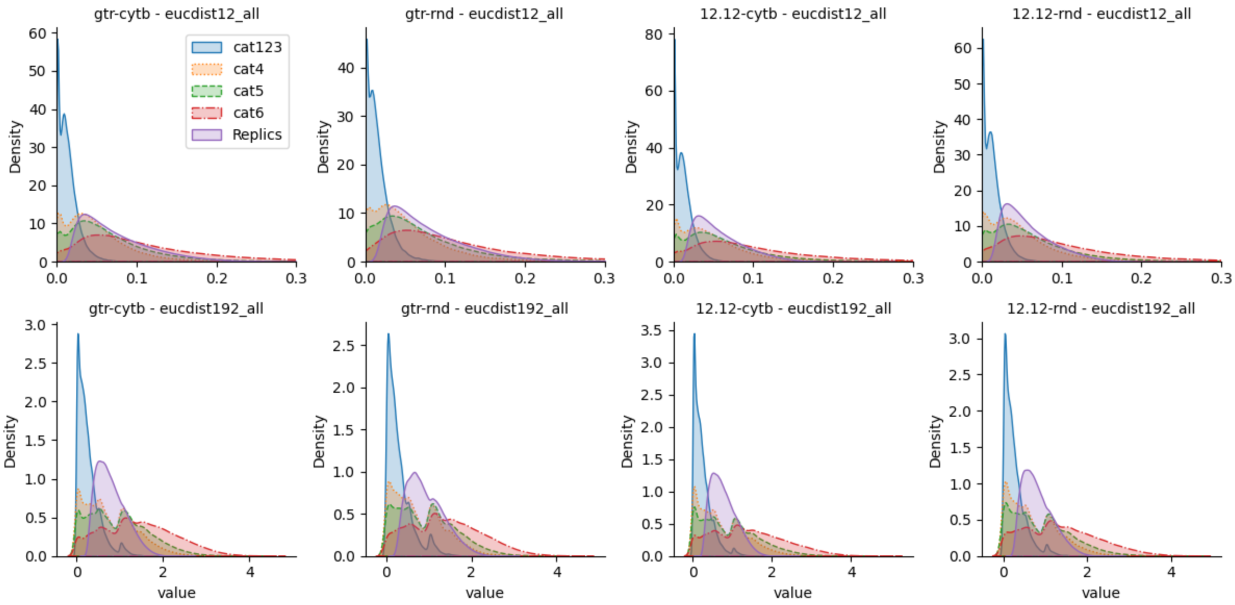


Fig. S8. Testing for ancestral states and/or mutational spectra computational accuracy on a real tree of 4705 mammalian species: the comparison between simulated and reconstructed 192 component (bottom row) and 12 component (top row) mutational spectra calculated using all mutations (synonymous and non-synonymous) based on Euclidean distances.

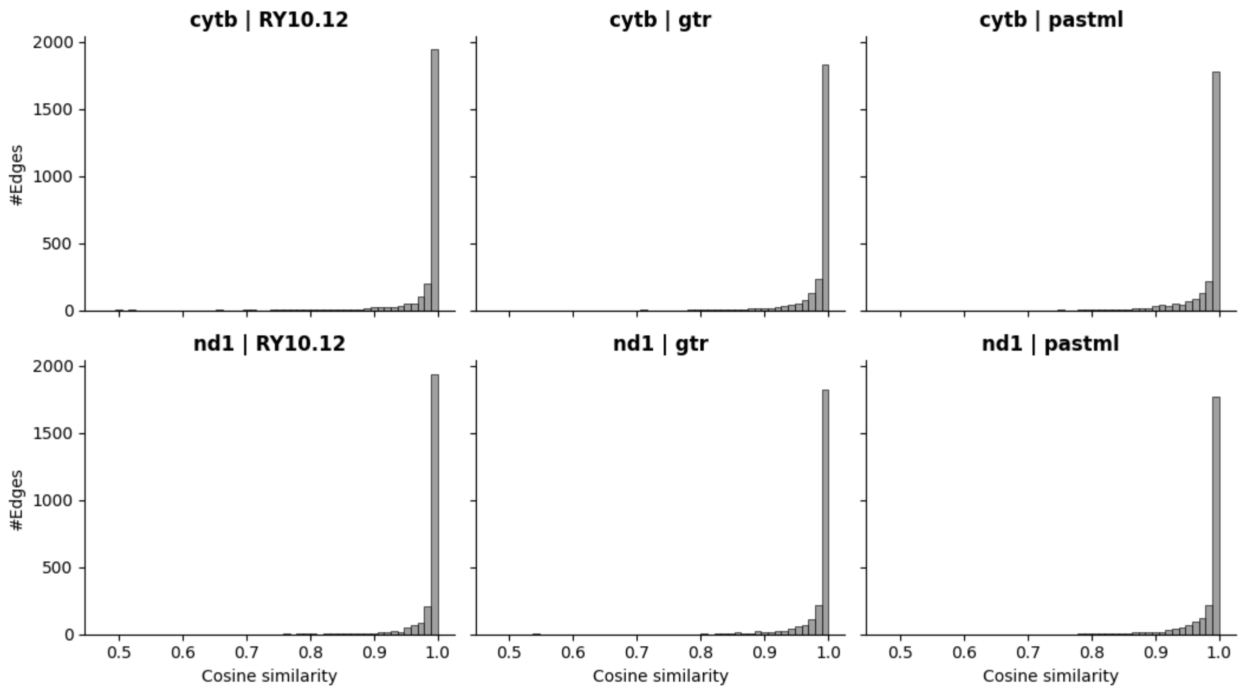


Fig. S9. The distributions (for each tree branch) of pairwise cosine similarities between two mutation spectra normalisations : by the frequency of trinucleotides, in a COSMIC - like way, and by the frequency of trinucleotides from all possible synonymous alternatives, in Nei and Gojobori - like way.

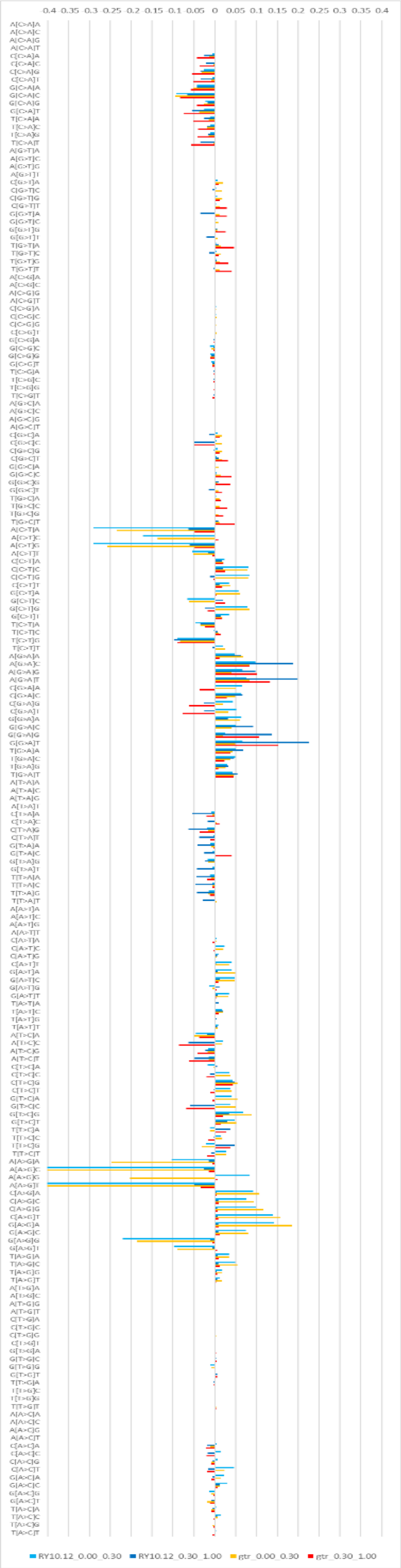
-0.4 -0.35 -0.3 -0.25 -0.2 -0.15 -0.1 -0.05 0 0.05 0.1 0.15 0.2 0.25 0.3 0.35 0.4
RY10.12_0.00_0.30
RY10.12_0.30_1.00
gtr_0.00_0.30
gtr_0.30_1.00

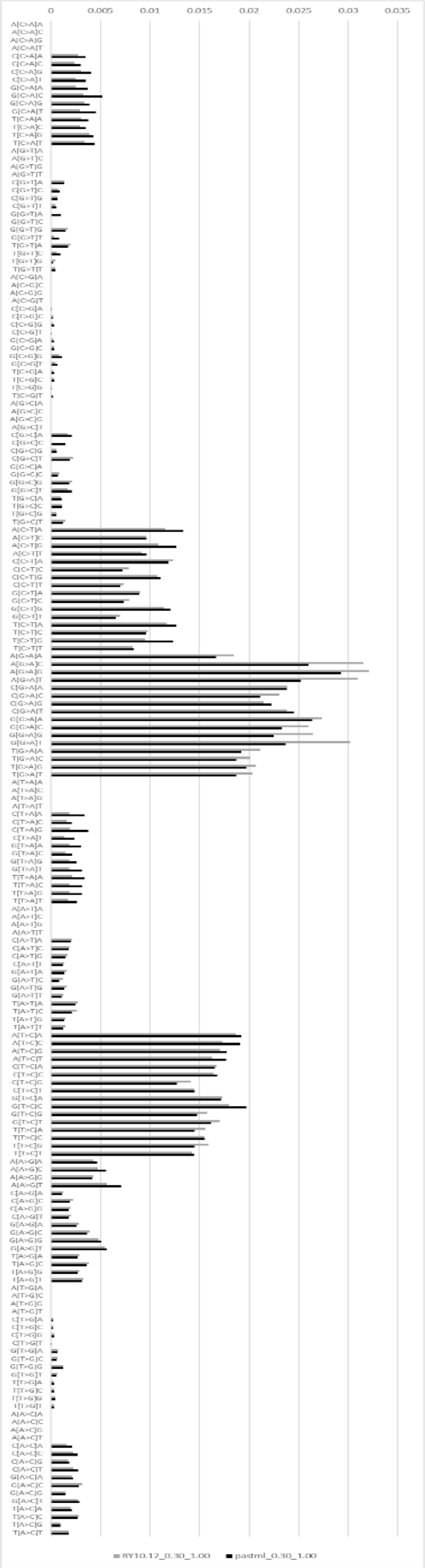
0 0.005 0.01 0.015 0.02 0.025 0.03 0.035
RY10.12_0.30_1.00
pastml_0.30_1.00

Fig. S10. The Mammals CYTB mutation spectrum, RateCat [5, 6] used. Left panel, the comparison of spectra obtained from PASTML-reconstructed ancestors with our approach: red and orange bars correspond to GTR substitution model (red, $p>0.3$; orange, $p<0.3$), blue and cyan bars correspond to RY10.12 substitution model (blue, $p>0.3$; cyan, $p<0.3$). Right panel, the Mammals ND1 mutation spectra obtained by PASTML (black) and by our approach, RY10.12 substitution model (gray); both our and PASTML reconstructions based on $p>0.3$ ancestral probability class.

SUPPLEMENTARY FILES

*Supplementary File 1. Additional species used in analysis of alignment sites variability.*
https://github.com/mitoclub/nemu-pipeline/blob/master/data/share/hominidae_and_mus_species.txt

*Supplementary File 2. Share of edges that have euclidean distances less than 5th percentile of reference distribution (replicas versus replicas).*
https://github.com/mitoclub/nemu-pipeline/blob/master/data/share/alisim_eucdist_mam_less_than_5percentile.csv

*Supplementary File 3. Share of edges that have euclidean distances greater than 95th percentile of reference distribution (replicas versus replicas).*
https://github.com/mitoclub/nemu-pipeline/blob/master/data/share/alisim_eucdist_mam_more_than_95percentile.csv
